# Full epistatic interaction maps retrieve part of missing heritability and improve phenotypic prediction

**DOI:** 10.1101/2022.07.20.500572

**Authors:** Clément Carré, Jean Baptiste Carluer, Christian Chaux, Nicolas Roche, André Mas, Gabriel Krouk

## Abstract

The first Genome Wide Association Studies (GWAS) shed light on the concept of missing heritability. It constitutes a mystery with transcending consequences from plant to human genetics. This mystery lies in the fact that a large proportion of phenotypes are not explained by unique or simple genomic modifications. One has to invoke genetic interactions among different loci, also known as epistasis, to partly account for it. However, current GWAS statistical models are moderately scalable, very sensitive to False Discovery Rate (FDR) corrections and, even combined with High Performance Computing (HPC), they can take years to evaluate for a full combinatorial epistatic space for a single phenotype. Here we propose a modeling approach, named Next-Gen GWAS (NGG) that evaluates, within hours, >60 billions of single nucleotide polymorphism (SNP) combinatorial first-order interactions, on a reasonable computer power. We first benchmark NGG on state of the art GWAS model results, and applied this to *Arabidopsis thaliana* providing 2D epistatic maps at gene resolution. We demonstrate on several phenotypes that a large proportion of the missing heritability can i) be retrieved with this modeling approach, ii) indeed lies in epistatic interactions and iii) can be used to improve phenotype prediction.

---

During the past decade, genome-wide association studies (GWAS) have allowed the discovery of many genetic variants associated with human (*1*–*3*), plant (*4*), animal (*5*) phenotypic traits. GWAS success is thus a reality and many discoveries made with this technique led to disruptive insights in biology, impacting basic knowledge as well as translational approaches to agronomy and medicine (*6*). However, when we observe on the one hand the striking resemblance of human twins, and on the other hand the amount of variation explained by GWAS signals, we are inclined to admit that “mono-dimensional GWAS” (study of genetic variation effect taken one at a time) is somehow limited. The missing heritability (*7,8*) can at least partially be explained by variant interactions, ie: epistasis. Approaching epistasis is a difficult problem that lies in the fact that current mathematical models linking genetic variations to phenotypes are extremely sensitive to up-scaling (in particular to the number of individuals in the study) and to False Discovery Rate corrections (*9*).

To attempt to make full epistatic maps a reality, we decided to use a radically different mathematical formalism combined with solving systems meant to take advantage of Graphic Processing Units (GPUs) being increasingly popular thanks to the rise of gaming and deep learning (*10*). For this, we established the NGG model that states and define heritability in this framework as done before by Zuk et al. (*11*) and others: We define *X* the matrix with n rows and p columns containing the genetic information. Each column displays the coded genetic variants (SNP) for the n individuals. We also define *Y* a vector containing the phenotype. The broad-sense heritability *H* may be defined via the following nonparametric “random signal plus noise” model : *Y*= *f*(*X*) + ε (NP) where the function *f* is unknown and general and ε is a random noise, independent from *X*that collects all other effects (other than genetic) on the phenotype *Y*, such as environmental effects for instance. Thus, the broad-sense heritability is expressed as *H* = *var*(*f*(*X*))/*var*(ε). The narrow-sense heritability *h* also sometimes named additive heritability accounts for part of the variance explained by genetics in the linear model *Y*= *X*θ + ε (L). The definition is *h* = *var*(*X*θ)/*var*(ε). We note that, of course, *model*(*L*) ⊂ *model*(*NP*). Notice for further use that since the slope parameter θ is unknown, the narrow sense heritability cannot be computed but only estimated (for example, via a plug-in estimator 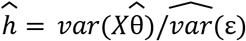. At last when the estimation method is Ordinary Least Squares (OLS) or some of its (regularized/penalized) variants, the definition above matches the classical *R* ^2^ and adjusted *R*^2^. Below the adjusted *R*^2^ is preferred for reasons related to both the dimensionality of the data (usually p is much larger than n) and the well-known inflation of *R* ^2^.

We further consider two models: Model1: *Y*= *X*θ + ε

Model 2 : *Y*= *X*θ_1_ + *Z*θ^2^ +_1_ ε

Where *Z* = *X*⋆ *X*is the partial face-splitting (or transposed Khatri Rao product) of matrices

(*12*). For self-containedness notice that when : 0

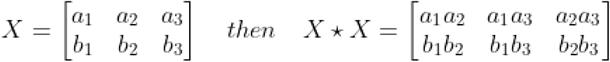

Matrix *Z* contains all the pairwise Kronecker products of columns of *X*, excluding the products of a column with itself. This matrix *Z* will be referred to as the matrix of interactions or shortly 2D-matrix. When *X*has p columns, *Z* has p(p-1)/2 columns. The matrix Z captures all interactions between the SNP’s. Although Model 2 remains linear, it is not additive anymore and bridges between Model L (or Model 1) and Model NP. Our resolution method employs acceleration techniques for regularized least square estimation in a sparse linear model (see Methods). These recent techniques from machine learning are coupled with a specific HPC architecture: GPU (See Supplementary Material). The outcome is a sparse estimate 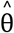 collecting the effects of each variant and each SNP interaction, instead of retrieving p-values as a regular GWAS does. As such, NGG can be seen as a sparse signal detection analysis and does not use multiple statistical testing which preclude the use of FDR correction. This classical correction is replaced here by a drastic procedure for variable selection.

To evaluate the performance of our model to retrieve epistatic signals, we first worked on simulated data. The simulation has been performed in two steps. First, we simulated *X*and *Y*(Fig.1A and 1B), second we simulated *Y* for real *X* (SNP matrix) retrieved from the 1001 genome project (*13*) (Fig.1C and 1D, Supplementary Material S1). These simulations are built to control narrow sense heritability (h^2^) of the trait (Fig.1). Using simulations, we show that the NGG formalism is able to capture simulated epistatic events for a wide range of model parametric values (see Supplementary Material S1). We found that Model 2 is quite resilient to noise but sensitive to the number of individuals used for the analysis (as discussed further, see remarks on Very High Dimension), as it radically improves for larger numbers of individuals (Supplementary Material S1).

**Fig. 1.**
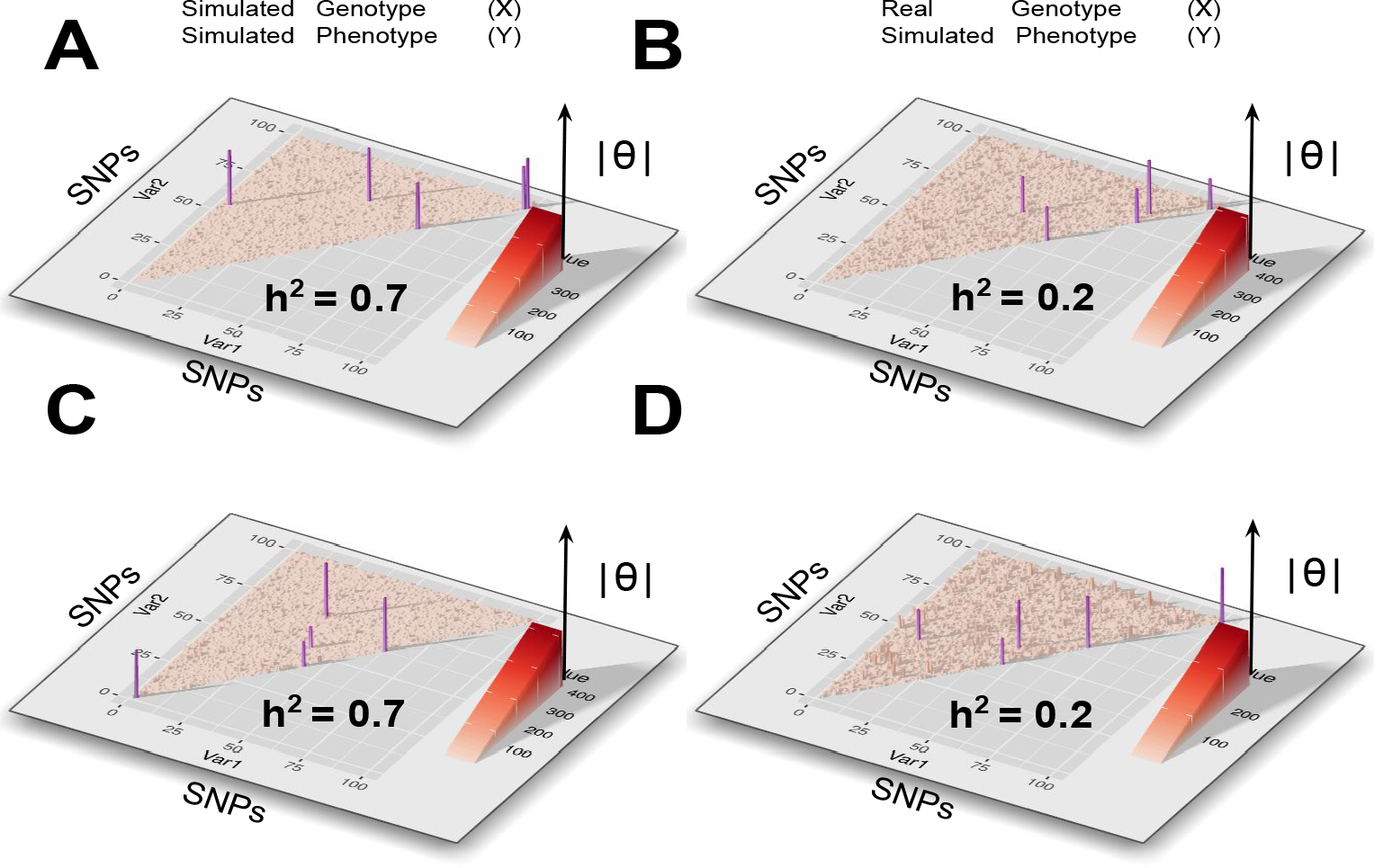
Next-Gen GWAS retrieves simulated epistatic interactions. Var1 (x axis) and Var2 (y axis) are a series of 100 SNPs. The triangle corresponds to SNPs combinations when the diagonal contains simple SNPs effects. z axis reports the estimated 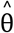 of simple SNPs (diagonal) and combinations (rest of the triangle). **A and B**) Genotype and phenotype data are simulated using specific and modulable parameters (see text for details on the simulation). Random noise is added. NGG retrieves the 5 simulated signals (purple) including the pure epistatic effects (outside of the diagonal). We have found that NGG is quite resilient to noise in the data (on phenotypes Sup info 1) and the power of NGG increases quickly with the number of individuals (Sup info 2). **C and D)** Phenotype data only have been simulated while genotypes are from the *Arabidopsis* genome (SNPs are sampled from X matrix). Again epistatic interactions (purple points) are retrieved by NGG.

We further benchmark our method on state-of-the-art available datasets and modeling approaches (*4, 14*). For this we first compared unidimensional (i.e. classical) GWAS results using the 107 Arabidopsis phenotypes studied in the landmark paper Atwell et al. [Ref. (*4*)]. We observed that major signals retrieved with EMMA algorithm (*4, 14*) are also retrieved by NGG (Fig. 2A and Supplementary Material S2 for the 107 phenotypes). For instance, EMMA and NGG methods both identify a major peak for the phenotype 88: *bacterial disease resistance* (Fig. 2A). This peak directly identifies the resistance gene RESISTANCE TO P. SYRINGAE PV MACULICOLA 1 (RPM1)(*15*). It is worth noting that for this particular phenotype, some signals emerge in NGG that are not detected by EMMA (Fig. 2) and that for certain phenotypes, NGG and EMMA converge towards a *x*^2^ relationship (Supplemental material S2). The opposite is also true although less frequent (see for the 107 phenotypes Supplementary Material S2). Furthermore, NGG clearly retrieves the effect of FLOWERING LOCUS C (FLC) a major gene in the control of flowering time (*16, 17*) in the top hits as compared to EMMA. Here we took this gene as an example for which NGG may be good at retrieving such important signals since it is intrinsically built to retrieve 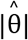, taking into account the other SNP effects (Model 1 and Model 2). For this we compared the capacity of NGG and EMMA to detect signals in the vicinity of the FLC locus (20 kb window). Interestingly Figure 2B shows that the NGG model indeed retrieves FLC as being the second strongest signal when EMMA reports it as the 30th signal (Fig. 2B).

**Fig. 2.**
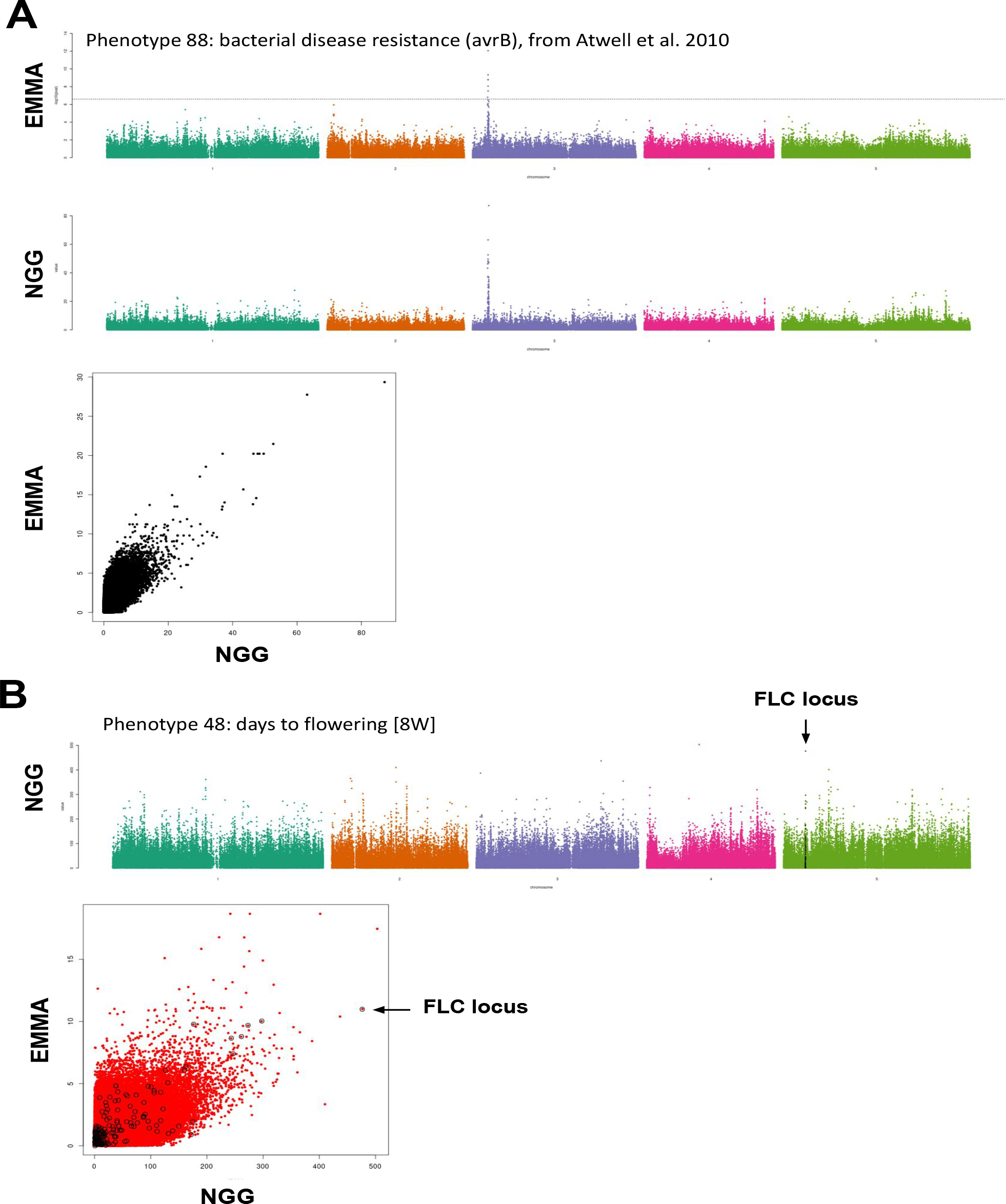
Next-Gen GWAS retrieves 1D-GWAS signal in Arabidopsis comparable to routinely used MML (EMMA (*4, 14, 23*)) and points to FLC locus for flowering phenotypes. A) Data from Atwell et al., (2010) have been used to compare efficiency of our algorithm to standards of GWAS in Arabidopsis. NGG and EMMA algorithms largely retrieve similar signals. B) The phenotype 48 (days to flowering trait [8W]) NGG results are displayed. SNPS in the close vicinity of FLC locus (a major component of flowering in plants) is represented by black dots. The scatter plot presents the fact that NGG better detects FLC effect as compared to EMMA.

We then compared the speed of our solution to the fastest GWAS to date, permGWAS (*18*). We evaluate that the computation time of one particular SNP effect or SNP combination effect (in Model 1 or 2 respectively) by our NGG algorithms takes around 4.4.10^−8^ seconds for 1000 samples or individuals. The GPU version of permGWAS, on a comparable (even more powerful) computer setup at ours, runs at a speed of 2.10^−4^ seconde per SNP. These four orders of magnitude differences bring the calculation of a full epistatic landscape by NGG below an hour for ∼60 billions SNP or combination of SNPs making full epistatic maps possible.

Being confident that NGG has the potential to point to true epistatic effects (Fig. 1) and having in mind that the number of individuals greatly improves the detection capacity of our model (Sup. Info 1), we took advantage of the recent work by Campos et al. 2021 (*19*) which is, to our knowledge, one of the available phenotype dataset with the greater number of Arabidopsis ecotypes to date. In this work Campos et al. (2021) provide the elementary composition (18 different elements) for >1100 different Arabidopsis ecotypes having been fully sequenced by the *1001 genome project* (*13*). Figure 3 reports results of unidimensional NGG for Phosphorus content (noted P31) of Arabidopsis leaves, that can be displayed at the same time as i) *support for the model x* |*SNP effect*|, or as ii) pure *SNP effect* (θ). The latter provides a new kind of Manhattan plot with negative values that can be interpreted as the SNP having a negative effect on the phenotype as compared to the reference genome (here *Columbia-0* ecotype) (Fig.3). Also, the effect reported in this new kind of Manhattan plot is now expected to be directly proportional to the effect of the genetic variation as compared to Col-0 phenotype, helping to choose for the best variant or gene to study.

**Fig. 3.**
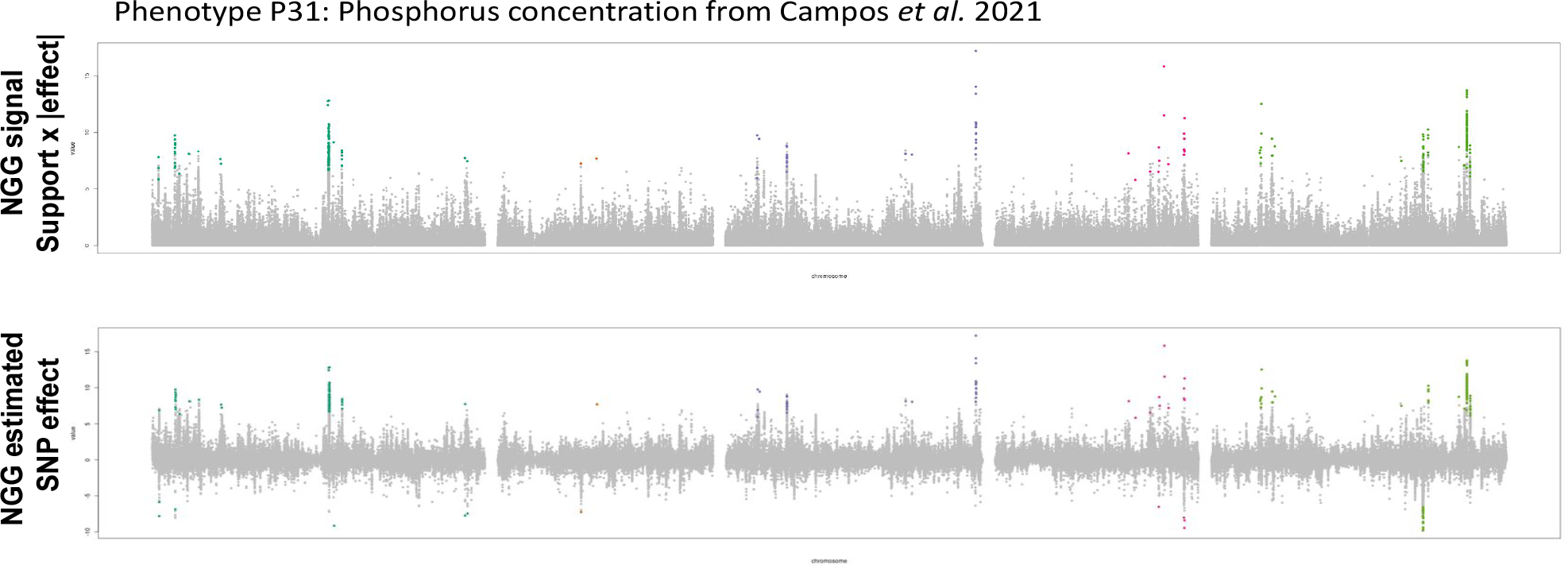
NGG provides direct estimation of SNP effect (θ) on the phenotype (Col-0 being the reference genome). The upper plot presents the NGG signal combining support (effect or not) x absolute value of the estimated effect of the genetic variation. The lower plot reports the estimated effect of each SNP. Colored data points are emerging from the noise in a bootstrap procedure described in Supplementary Material 1).

We further proceeded with the computation of full epistatic maps or 2D-NGG for phenotypes retrieved from the Campos et al. (2021) and Atwell et al (2010) datasets. We focused on these dataset as they present a relatively high number of ecotypes (>1000), and an important diversity of well known phenotypes respectively. To do this, we prefiltered SNPs having a particular Minor Allele Frequency (MAF) because the probability for the combination of SNPs (that we call MIAF for Minor Interaction Allele Frequency) to be of interest for epistatic measurements directly depends on the MAF as *Z* = *X*⋆ *X*(above). For this we prefiltered 346094 SNPs, for Campos et al, and between 341067 and 371956 SNPs, for Atwell et al, having a MAF greater than 0.3. The full epistatic landscape is thus 59.890 billion interactions for Campos et al. (2021) and between 58.163 and 69.175 billion interactions for Atwell et al. (2010).

Nowadays, this quantity of data represents a challenge on its own to compute, store, and display the results as it relates to a “Very High-Dimensional” (VHD) framework (*20*). VHD is mathematically defined in terms of the size of the genotypic matrix *X* (*n* rows and *p* columns) and in terms of sparsity of the unknown parameter to be estimated or tested, here the number of “active” SNPs and interactions for a given phenotype: *k*. In this framework (*20*), we can evaluate the effects of VHD genotypic input matrices on the performance of several popular methodologies (for hypothesis testing, support estimation, prediction) and shows that when *k log(p/k)* is large with respect to *n* then statistical estimation and testing errors inflates dramatically. We believe that this is at least in part the reason for which full epistatic maps (2D-GWAS) were so far out of reach.

Following this line, in our study (Fig. 4) n=999, and p is around 60 billions. Following a reasonable estimate for sparsity *k*, granting satisfactory and reliable outputs, should be not more than 50. This is why, in our forthcoming study we mainly consider and analyze in a final stage around 30 significant SNPs interactions or composite components.

**Fig. 4.**
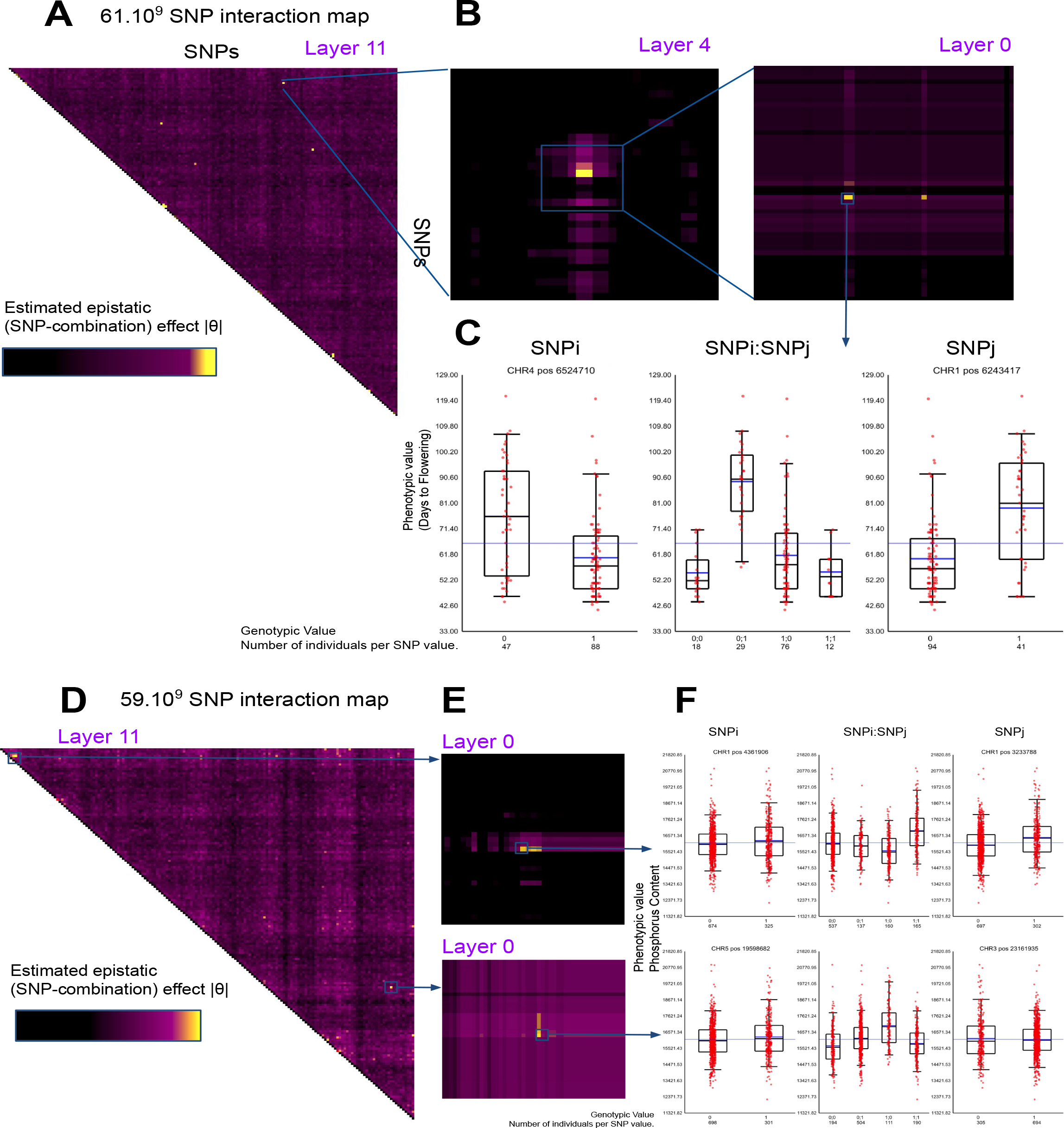
2D-NGG results provide an estimation of 61.2 billion SNP combination effect. for: A) Atwell et al, phenotype ID:31, days to flowering time FT10, B) Phosphorus content Campos et al. 2021 measured by ICP-MS. The results are presented as heatmaps and histograms to observe the epistatic interactions between SNPs.

Figure 4 displays 2D-NGG results for i) Arabidopsis flowering times (Fig. 4A)(*4*), ii) Arabidopsis phosphorus (P31) leaf content (Fig. 4B) (*19*). Results are displayed as a square heatmap triangle for which ∼60 billion signals 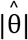 are provided. One 2D-NGG result dataset represents ∼500 Go of data. To navigate and mind this large dataset a visualization tool has been developed named *Luciol* that can be understood as a “Google Earth” for full epistatic maps. In short, results are organized in layers as such the max intensity of a genomic region is reported on the higher layers. Here in Figure 4, layer 11 represents our maximum zoom-out condition. A zoom between layer 11 to layer 0 (layer for which a given pixel represents a direct SNP/SNP combination) corresponds to a 4.2 million times zoom. In other words a pixel in layer 11 reports the max intensity of 4.2 million real SNP/SNP interactions underlying layer 0. Observation of full epistatic maps as well as local signal informs on the genetic architecture underlying a given phenotype (Fig. 4).

In the case of the flowering trait (Fig. 4A), around 6 major epistatic signals emerge where 2 of them are close to the diagonal. The proximity of the diagonal refers to potential epistatic interactions of neighboring genes (although few Mb away). To display unambiguous epistasis we thus decided to report here the fourth stronger effect that lies very far from the diagonal. A zoom at the 2D-locus reveals the structure of a 2D-GWAS peak that appears bi-modal (ie: supported by at least 2 distant SNP combinations, 2 local bright spots in the epistatic map, Fig. 4B). This peak points to 2 loci predicted to be epistatic. The first loci is at position CHR4:6524710, and the second one is at CHR1:6243417. Using these coordinates, the matrix X and phenotype Y are parsed to plot the phenotypic distribution following the combination of SNPs (a sort of 2D-haplogroup). This is reported by the box plot in Figure 4C. Herein we observe that this epistatic effect involves 2 loci having a moderate effect individually as reported to the SNPi and SNPj boxplots (left and right panels). However, the combination of the simple effect cannot predict the effect of the combination since the positive effect of SNPj, from 0 to 1 modality, seems enhanced by the SNPi (0) modality and totally repressed by the SNPi (1) modality. This clearly indicates the potential presence of an epistatic effect between these 2 loci. We also report (Fig. 4D to 4F) the epistatic interactions in the control of plant leaf phosphorus content. This epistatic map reports around 8 strong epistatic signals. Here we zoom into 2 of them being the stronger ones with respect to their predicted value (|θ|).The first one is relatively close to the diagonal although both epistatic SNPs lie in chromosome 1 but eleven Mbp away (Fig. 4F top panel). The second one concerns an epistatic effect predicted to involve SNPs on 2 different chromosomes namely CHR5 and CHR3 (Fig. 4F bottom panel). These 2 epistatic signals are built upon the effect of a strong combination of SNP effects as it appears impossible to predict the combinatorial output of these SNPs by solely analyzing the effect of the simple SNP modalities (compare box plots of SNPi and SNPj to box plot of SNPi:SNPj). Here, these effects can totally be missed by previous studies of epistasis that, to date, necessarily implied a selection of genetic variables (*21*).

We then decided to evaluate the quantity of heritability (*h* ^2^) retrieved from 2D-GWAS as compared to regular 1D-GWAS (Fig. 5). The differential heritability between 1D and 1D+2D GWAS was estimated by Principal Component Regression (PCR) (*22*), carried out on a set of selected SNP and SNP-interactions (Fig 5A). The principle of PCR dates back to the late 50’s (*22*). PCR combines Principal Component Analysis (PCA) on the input features of a model followed by linear regression (*22*). First a PCA of X provides a low number of principal components and a dimension reduction by selecting fewer components associated with the highest eigenvalues modulus of X. Regression is then performed on this reduced set of components (related to the VHD problem that we described above) that play the role of new synthetic inputs. PCA concentrates the information of the large matrix X or Z in a smaller matrix, removing collinearity as well because the components are, by definition, not correlated.

**Fig. 5.**
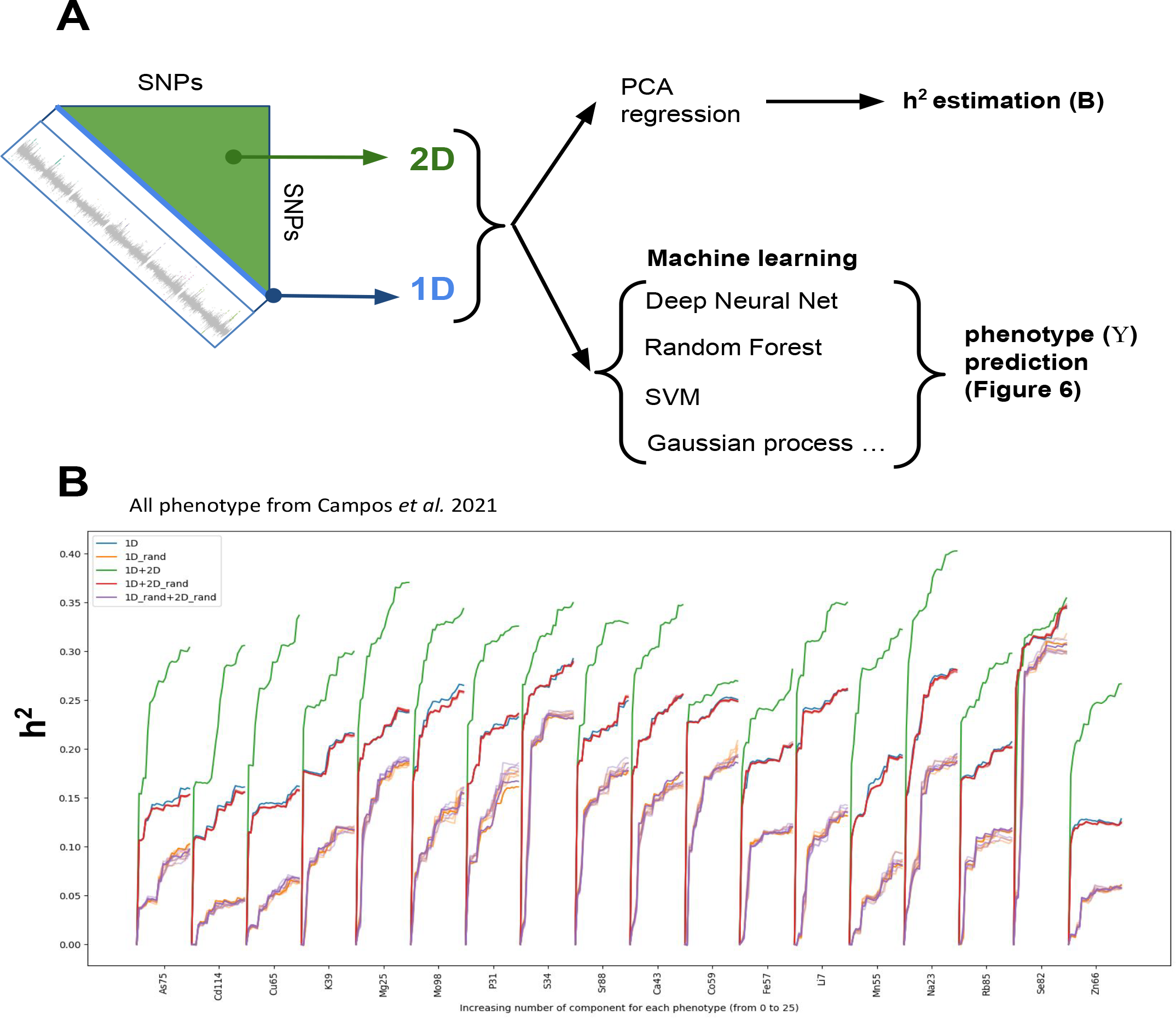
Estimation of retrieved missing heritability. A) Analysis scheme employed to estimate retrieved heritability and phenotypic predictions (in Fig. 6) from 1D signals (Blue Diagonal), and from 2D NGG signals orange triangle representing 61 billion interactions. B) Heritability (h^2^ seen as adjusted R^2^) is measured for an increasing number of PCA components, and for signal retrieved only from 1D-GWAS or 1D-GWAS (V data points) + 2D-NGG (W data points).

Here PCR is carried out i) on a set of p SNPs then, ii) on a set of the same SNPs as in (i) + q SNP/SNP-interactions (Fig. 5). The plots show the retrieved “missing heritability” (difference between the green line and blue [1D signal] or red lines [1D signal plus 2D_random]) as measured by the adjusted R^2^ when the number of selected components increases (x axis Fig. 5B). For the vast majority (16 of the 18 phenotypes) a good proportion of heritability is retrieved in the 2D signals. Only, Cobalt or Selenium do not display a radical improvement in the explained variance. By applying this method we observe that information in the epistatic landscape indeed contains a good proportion of the missing heritability (Fig. 5B). For the Phosphorus content of Arabidospis leaves for instance the heritability measures in the 1D GWAS ranges around 22%. h^2^ then increases to 33% when the information in the 2D-GWAS is considered.

Having at hand a view of the full epistatic maps can be seen as a new route towards at least two kinds of developments. The first one is obviously experimental validations. These are very labor intensive and may require a very long time to precisely dissect epistatic interactions. These are under investigation but we wished to release our results in the light of the second. The second one is phenotypic prediction. Indeed, one could see the NGG as a very strong variable selection process (Fig 5A) that may benefit precision medicine or several programs of agronomic selection for plants or animals. We thus further evaluate the role of NGG signals for phenotypic predictions through the use of a broad set of proper machine learning algorithms including Deep Neural Networks (DNN), Support Vector Machine (SVM), Gaussian Processes (GP), Gradient boosting (GB), Random Forest (RF), Linear regression, Lasso, Elastic Net. These techniques were used to predict the 18 phenotypes from the Campos et al (2021) work (described above). We also crossed these machine learning techniques with an increasing number of 1D or 2D signals/SNPs (50, 100, 500, 1000, 5000, 10000). To perform a proper control we repeated this *in silico* experiment but instead of providing the models with proper 2D signals, we randomly sampled epistatic signals (named 2D-random) to evaluate our capacity to predict plant mineral content. As classification problems are easier to solve and that the number of individuals is still a bit limited for regression approaches, we also used quantiles to rank phenotypes into 5 or 3 classes. By crossing all these parameters, we ended up with 1728 different models for 1D+2D signals (y axis of the plot Fig. 6A) and the same number of models for 1D+2D_random (x axis Fig.6A). Our capacity to predict phenotype is performed on 30% of the dataset (validation set) that were not used to fit or train the models. The quality of the models are evaluated through classical precision/recall curves and F scores. Figure 6 presents the F1 scores of best predicted classes (measure a precision/recall compromise) for 1D+2D random against 1D+2D models. All the models lying above the diagonal (x=y) correspond to models for which predictive power is improved by epistatic signals (Fig. 6A).

**Fig. 6.**
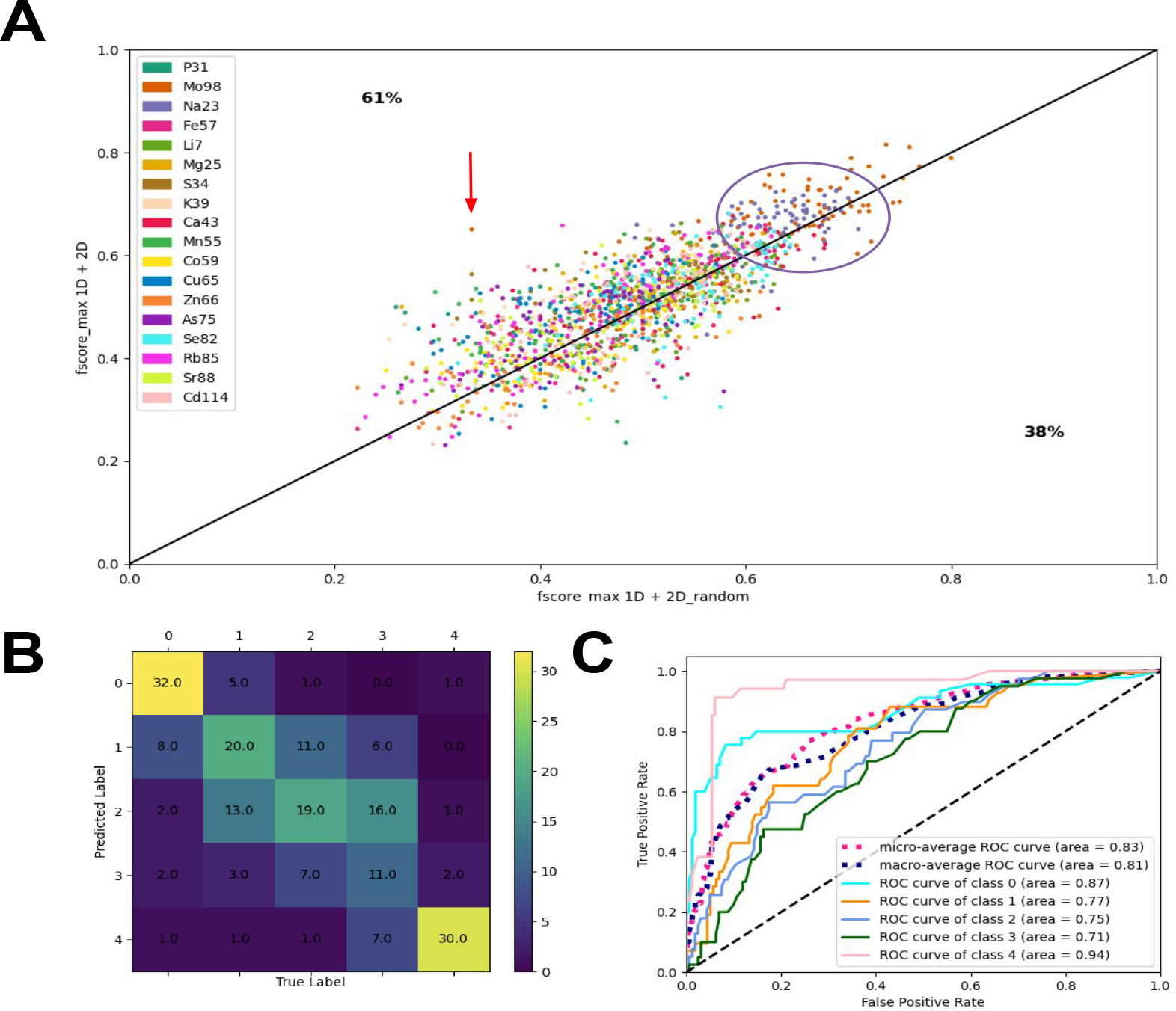
NGG retrieves genetic markers in epistatic signals improving machine learning procedures. A) In this dot plot each dot corresponds to a given machine learning model (among: SVM, RF, DNN, Gaussian processes, LASSO, Elastic-Net Classifier) trying to predict a given phenotype (18 elemental concentration of Arabidopsis leaves represented with different colors) combined with different learning data inputs including a different number of classes (3 or 5) and different number of SNPs (50, 100, 500, 1000, 5000, 10 000) (Full dataset provide in Sup Material 3). The x axis reports max F1 score for the mode provided with SNPs simple 1D signals and randomly picked 2D epistatic SNP combinations (our control). The y axis reports 1D signals and 2D signals retrieved by NGG for the sample model and parameter combinations, respectively. We observe a clear improvement (above the y=x line) of >65 % of the models. B,C) Example of the good prediction of the Molybdenum (Mo98 phenotype) classified concentrations. B) confusion matrix, C) panel ROC curves for each class of Mo98 phenotype.

We observe that 2D epistatic signals largely improve phenotypic classification (Fig. 6A) as 61% of the models are improved. Interestingly, models having a low F1 score and models having a high F1 score seem to beneficiate the most of the epistatic signals. We wish to highlight some particular points for which we observe a dramatic increase in our capacity to predict phenotypic classification (Fig. 6A). The red arrow Figure 6A points to a model for which 1D signal alone does not allow a good classification (value 0.33), when the same model with epistatic signal reaches a Fscore of 0.65. We also observe some phenotypes such as Na23 (sodium leaf content) for which most models and parametric values of the machine learning procedures globally beneficiate the epistatic signal showing that retrieved 2D signals are globally bringing new information (purple circle Fig. 6A). Figures 6B and 6C display an example of our capacity to predict phenotype classification for molybdenum leaf concentrations. This level of precision and recall opens avenues for plant selection procedures.

In conclusion, in this work, we provide a first dive into complete epistatic maps with enough SNP to reach a gene resolution and new tools to analyze this VHD problem. We apply it to the model plant *Arabidopsis thaliana* but our technique is fully generic and can easily reach other biological models. We demonstrate that, as hypothesized before (*11*), a substantial part of the missing heritability indeed lies in epistatic interactions (Fig. 5). We also show that this never observed 2D epistatic signal brings us a bit closer to the prediction of phenotypic values by machine learning procedures in plants, but we hope soon, in other biological models as well.

## METHODS

### Data

Arabidopsis dataset corresponds to data issued from the 1001 genome project (*13*) and kindly provided by Arthur Korte lab. It consists of a genotype matrix above mentioned as genotype or X matrix containing 9124892 SNPs and 1135 ecotypes. For NGG analysis MAF is controlled (0.3<MAF) resulting in a MAFed X’ matrix containing 346094 SNPs for Campos et al. (2021) and between 341067 and 371956 SNPs, for Atwell et al, (2010).

### Simulations

The simulations (Fig. 1) are performed on R. Code can be found at (https://github.com/CarluerJB)

### Computer power

This work has been performed on a PowerEdge T640 DELL Server, RAM 377 Go, 4 NVIDIA Quadro RTX 6000 (24Go).

## ACKNOWLEDGMENTS

We wish to thank Dr Sandrine Ruffel, Dr Benoit Lacombe and Dr Eric Hosy for their valuable remarks on the manuscript.

## Funding

Research on phenotype prediction is supported by the CNRS and by Montpellier University (Isite MUSE project AI3P) to G.K. and A.M. This work/project was publicly funded through ANR (the French National Research Agency) under the “Investissements d’avenir” programme with the reference ANR-16-IDEX-0006. The development of a first version of the computer algorithms was supported by SATT-AxLR and by Agropolis Fondation (SATT prematuration project) to G.K. A different version of the NGG resolution algorithm and the visualization algorithm named *Luciol* allowing data exploration has been developed by BionomeeX™.

## Author contributions

Conceptualization: C.C., G.K. and A.M. Methodology: C.C., J.B.C. (performed prediction, machine learning and simulations), G.K., A.M., N.R., Ch.Ch, Writing – original draft: C.C, J.B.C, A.M, G.K.

## Competing interests

C.C, G.K, A.M are co-founders of a CNRS and University of Montpellier spin-off company BionomeeX™ (BionomeeX.com) to provide a simple and rapid access to NGG technology and subsequent data analysis.

## Data and materials availability

Phenotypic and genotypic data have been retrieved online. All code associated with heritability and data modeling can be retrieved at GitHub (https://github.com/CarluerJB). Raw data results of 2D-NGG for the phenotypes presented Figure 4 are available upon request to any of the corresponding authors.

## Supplementary Materials

**Supplemental Figure 1. Influence of the number of individuals on the power of epistatic signal detection**.Var1 (x axis) and Var2 (y axis) are a series of 100 SNPs. The triangle corresponds to SNPs combinations when the diagonal contains simple SNPs effects. z axis reports the estimated θ of simple SNPs (diagonal) and combinations (rest of the triangle). Genotype and phenotype data are simulated using specific and modulable parameters (see Github for details and code). Random noise has been added. Simulated signals are in purple. Simulations control heritability (h^2^). The number of individuals (n) varies over the figures (see details above panel). The number of individuals has a substantial effect on GWAS signal detection.

**Supplemental Figure 2. Comparison of 100 GWAS results using EMMA or NGG algorithms**.Atwell et al. (2010) dataset has been analyzed using both algorithms. Results are provided as Manhattan plots and signals between EMMA and NGG are compared by plotting their respective signals against each other.

**Supplemental Table 1. Machine Learning Results**.This table reports scores (precision, recall, Fscores) for the prediction of 18 Arabidopsis Phenotypes (mineral content), and a combination of different parameters of the ML procedures, phenotypic classes, Machine Learning, using 1D signals or 2D signals as input variables. These data led to the making of Figure 6A.

