## Supplementary material for "Full epistatic interaction maps retrieve part of missing heritability and improve phenotypic prediction": Sup Fig 1

**Supplemental Figure 1.**

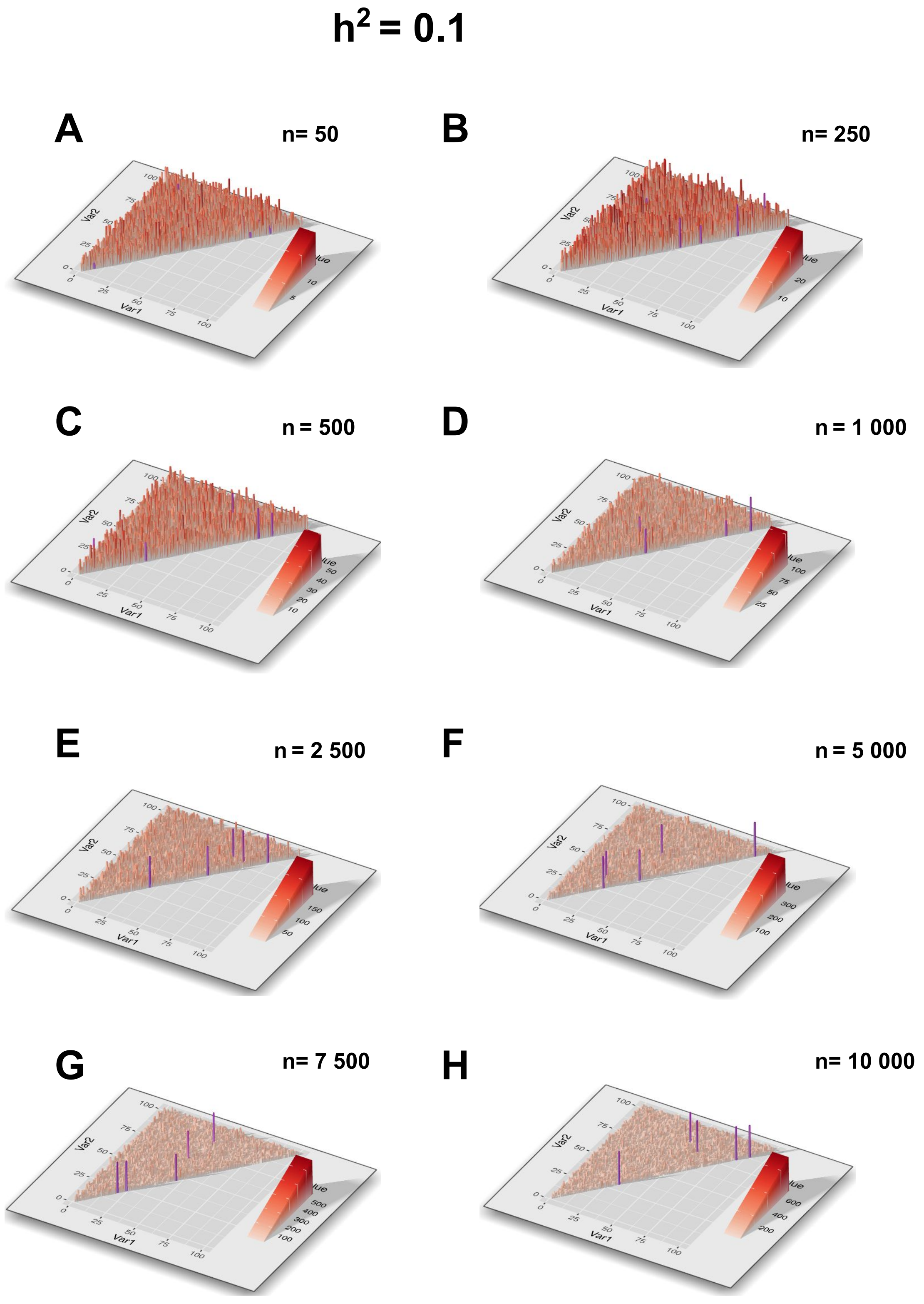


**Supplemental Figure 1. Influence of the number of individuals on the power of epistatic signal detection.**
Var1 (x axis) and Var2 (y axis) are a series of 100 SNPs. The triangle corresponds to SNPs combinations when the diagonal contains simple SNPs effects. z axis reports the estimated θ of simple SNPs (diagonal) and combinations (rest of the triangle). Genotype and phenotype data are simulated using specific and modulable parameters (see Github for details and code). Random noise has been added. Simulated signals are in purple. Simulations control heritability ($h^2$). The number of individuals (n) varies over the figures (see details above panel). The number of individuals has a substantial effect on GWAS signal detection.

**Supplemental Figure 1.**

# $h^2 = 0.2$

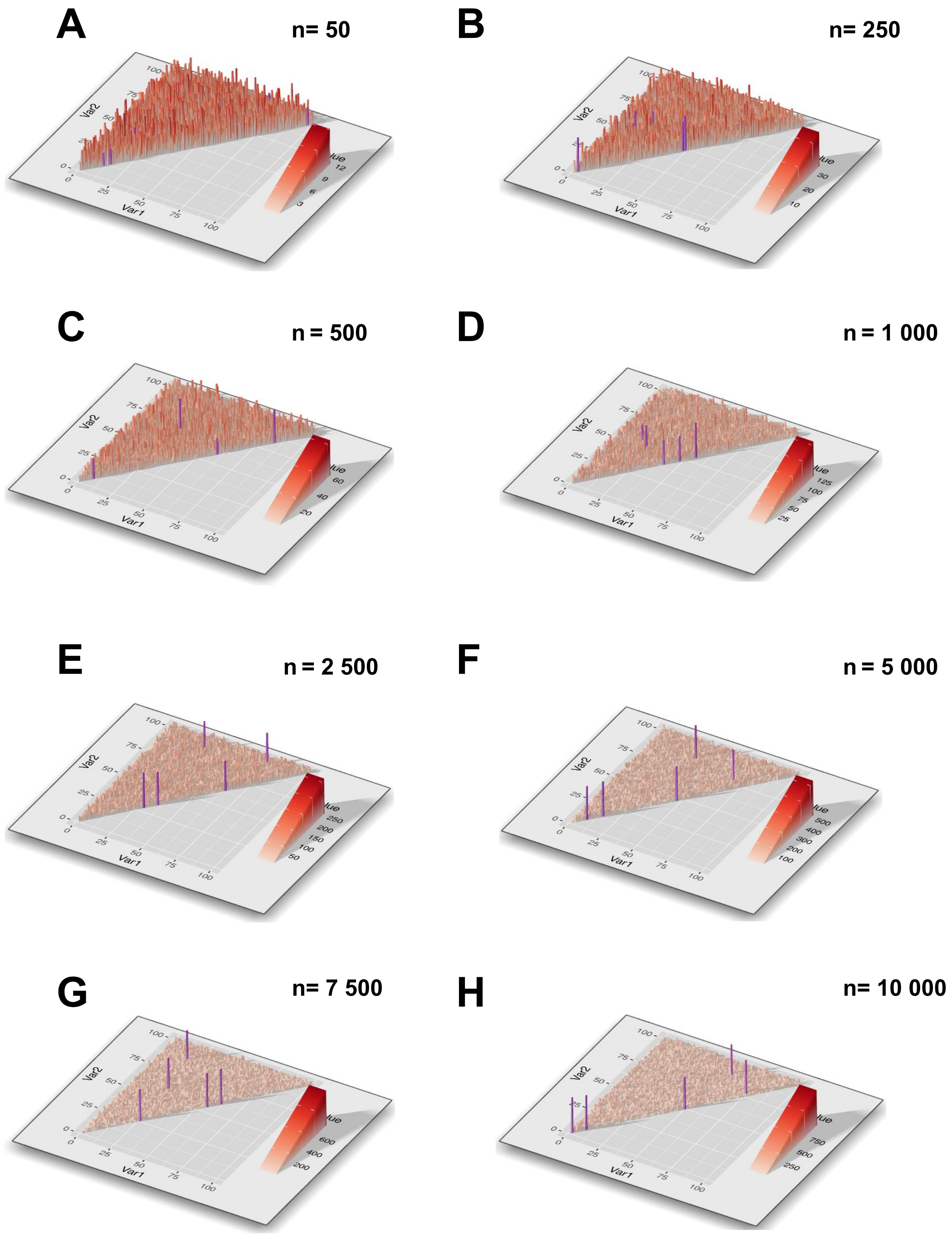

**Supplemental Figure 1.**

# $h^2 = 0.4$

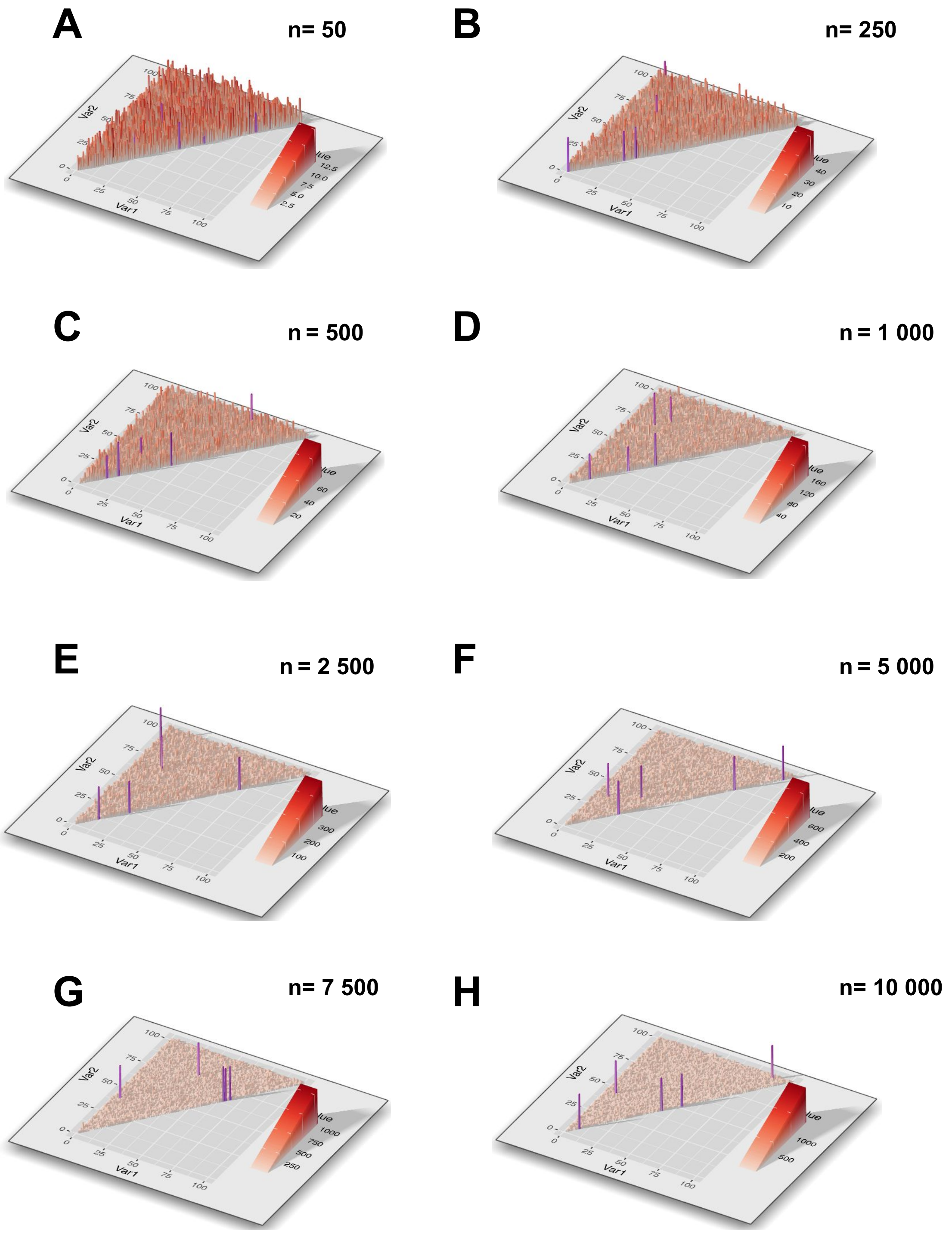

**Supplemental Figure 1.**

# $h^2 = 0.7$

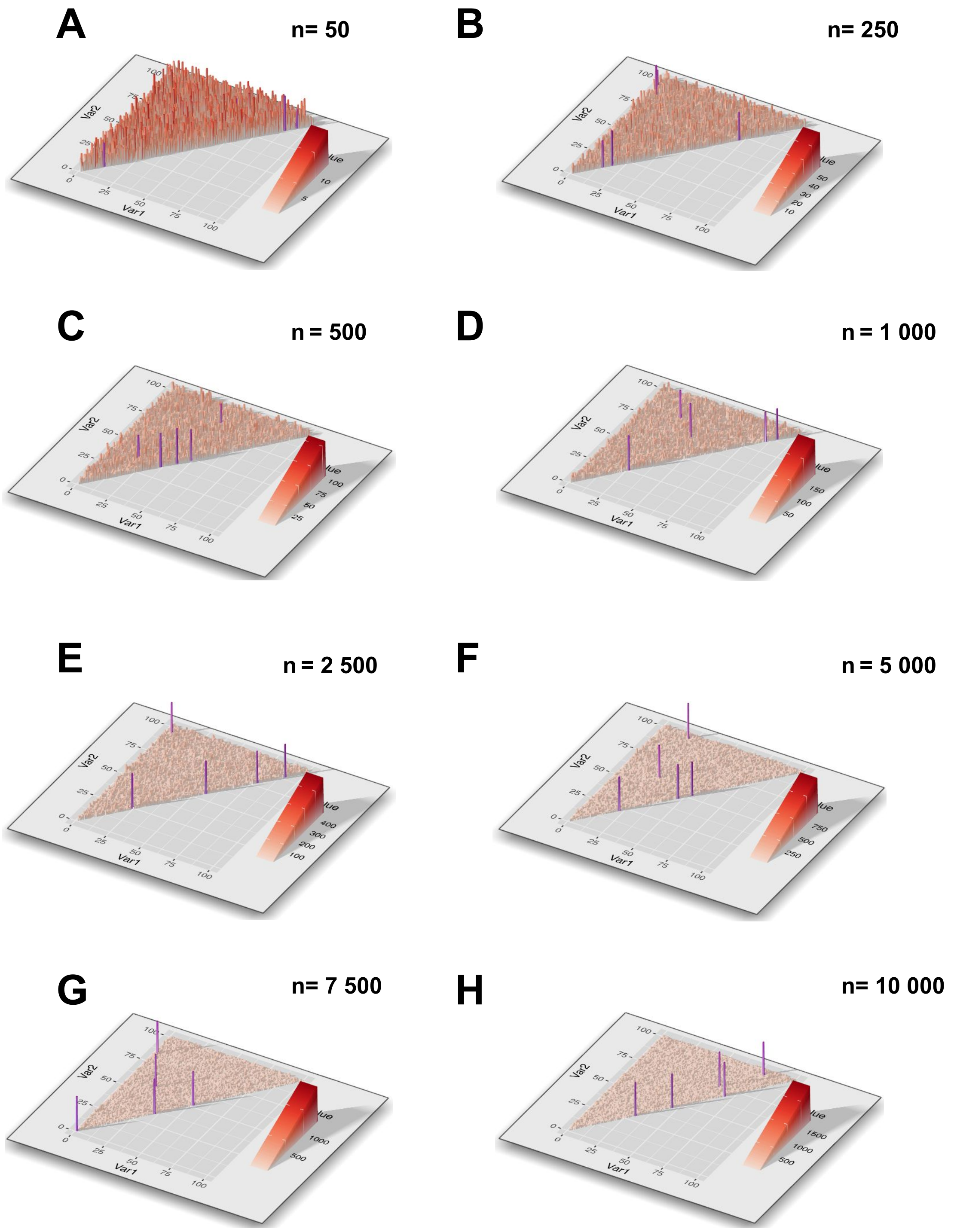
