## Supplementary material for "Full epistatic interaction maps retrieve part of missing heritability and improve phenotypic prediction": Sup Fig 2

1
leaf number
(FT Diameter Field)

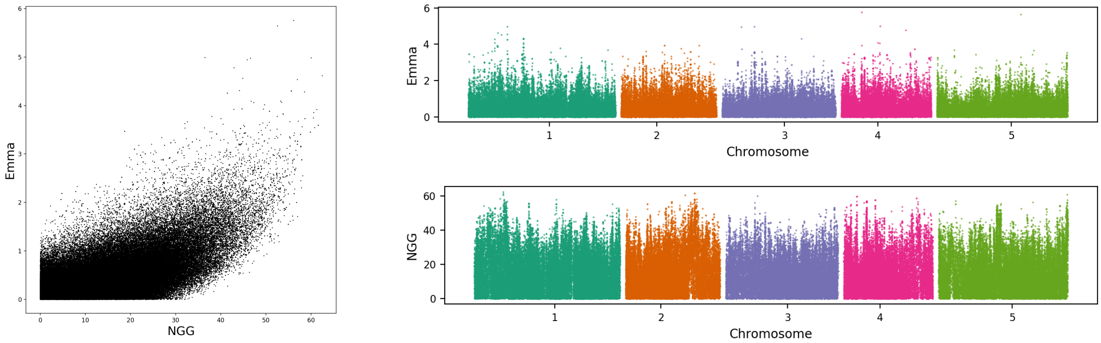


2
bacterial disease
resistance
(At2 CFU2)

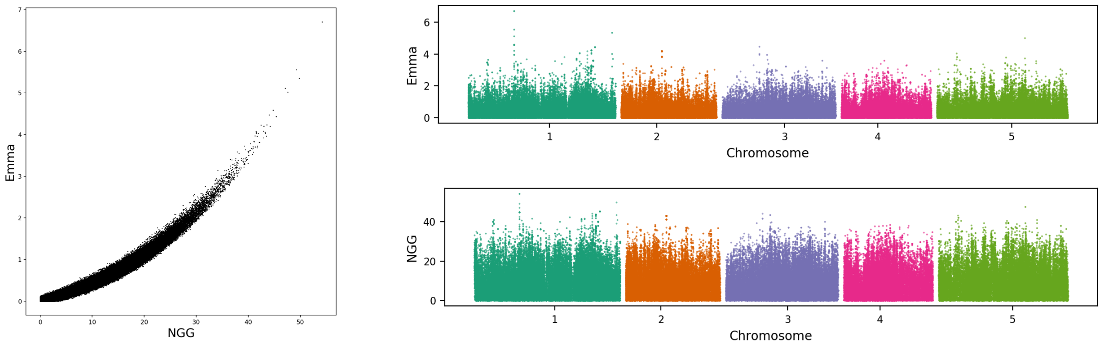


3
leaf margin serrated
(Leaf serr 16)

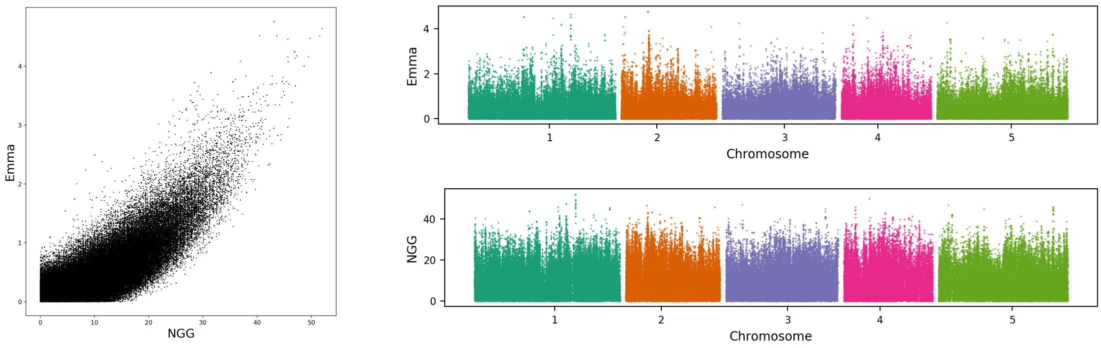


4
seed dormancy
(Seed bank 133-91)

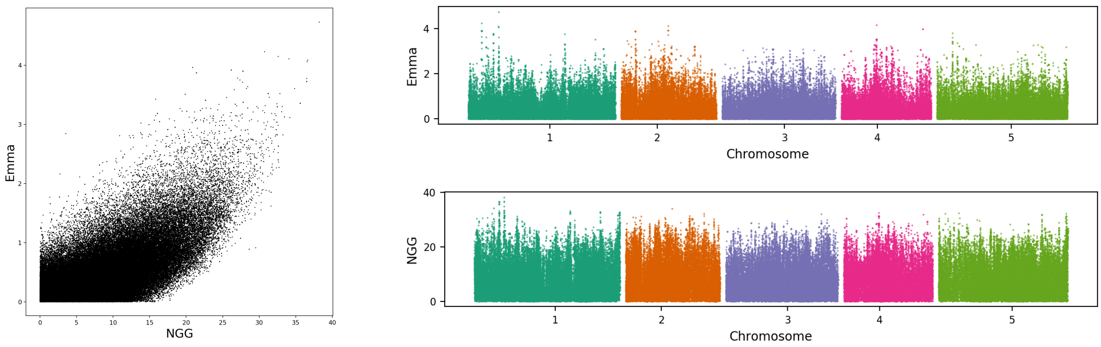

5
sodium
concentration
(Na23)

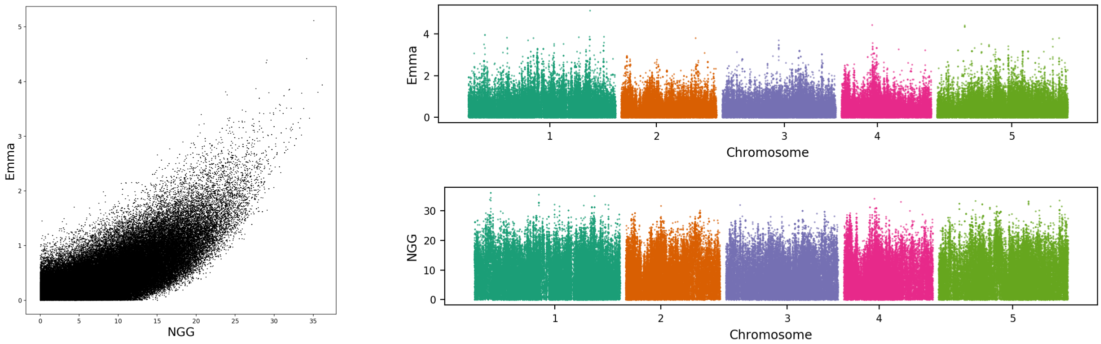


6
leaf margin serrated
(Leaf serr 10)

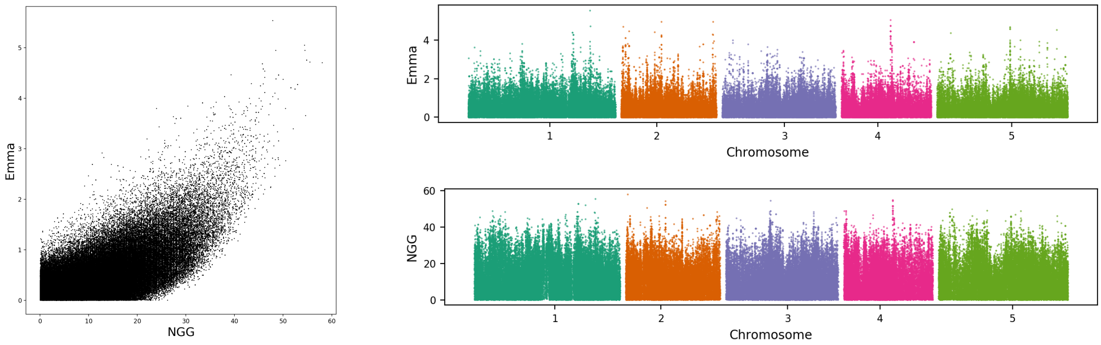


7
protist disease
resistance
(Emco5)

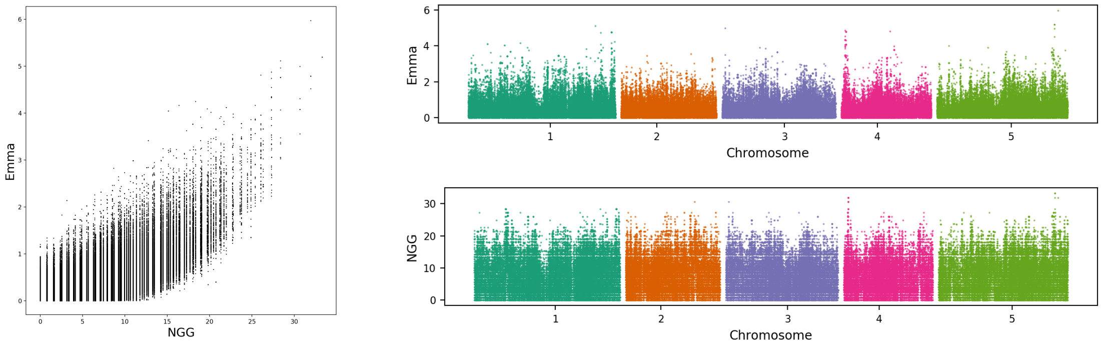


8
rolled leaf
(Leaf roll 16)

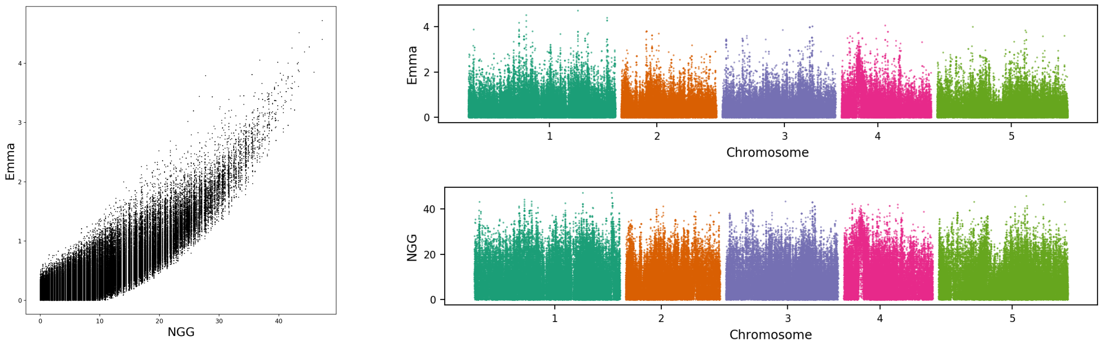

9
leaf margin serrated
(Leaf roll 10)

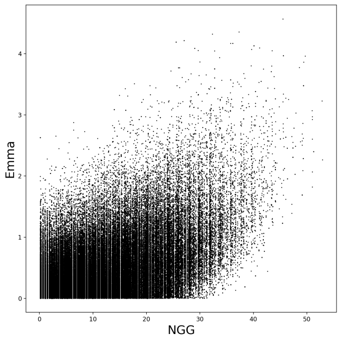


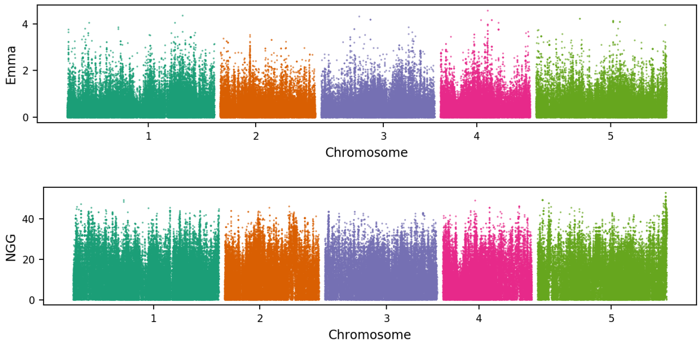


10
bacterial disease
resistance
(Bs)

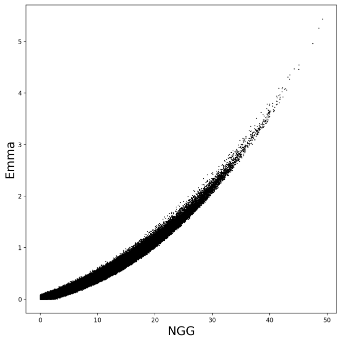


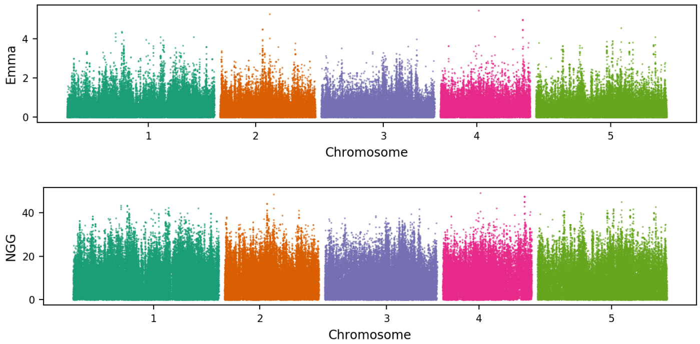


11
days to flowering
trait
(2W)

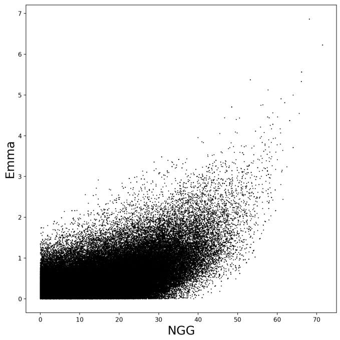


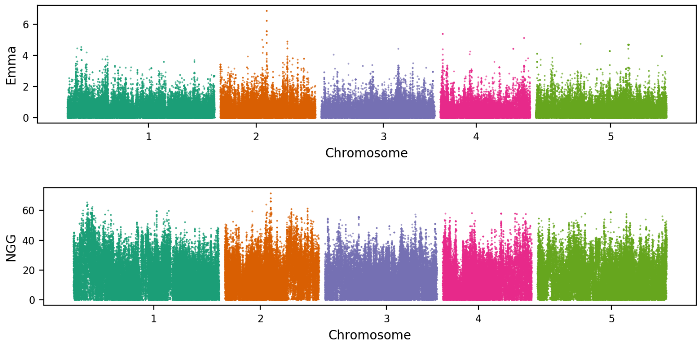


12
rolled leaf
(Rosette Erect 22)

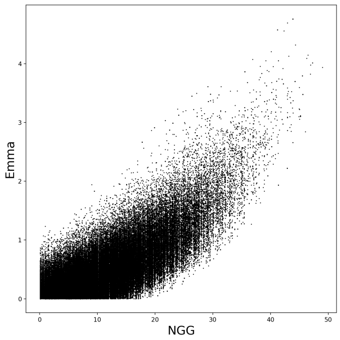


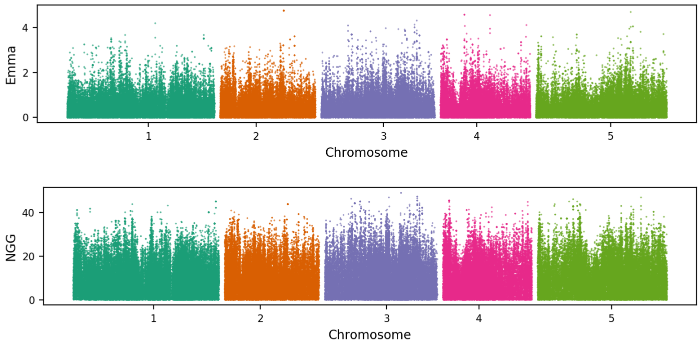

13
cadmium concentration (Cd114)

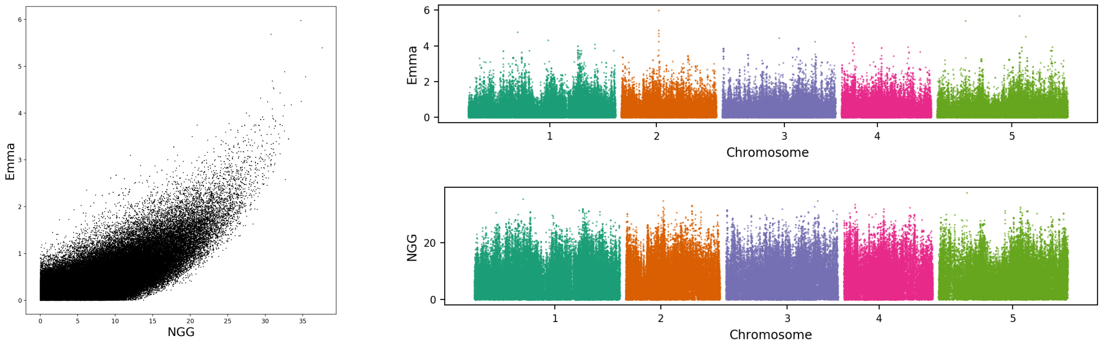


14
plant width (Width 16)

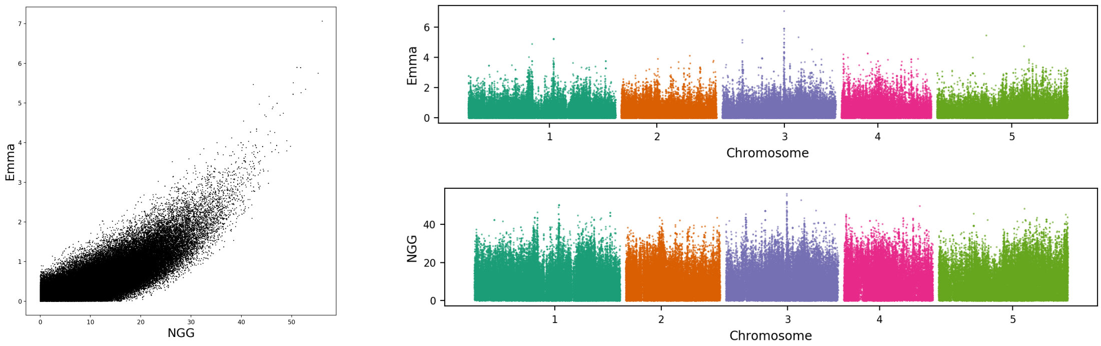


15
seed dormancy (Storage 28 days)

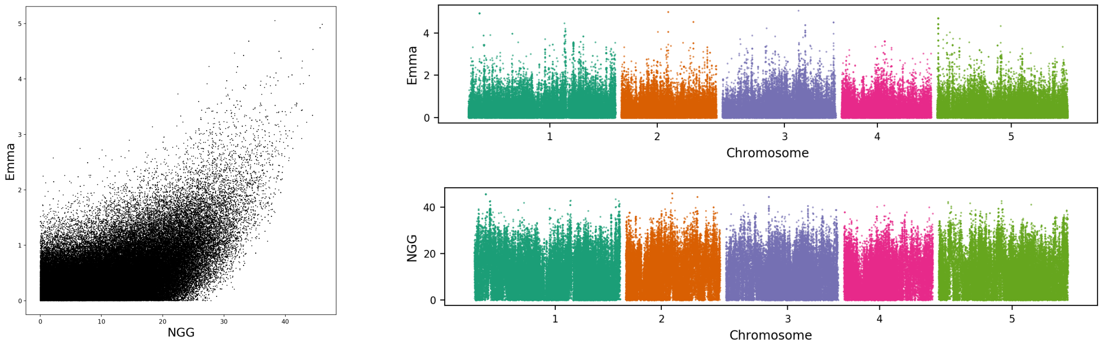


16
leaf necrosis (LY)

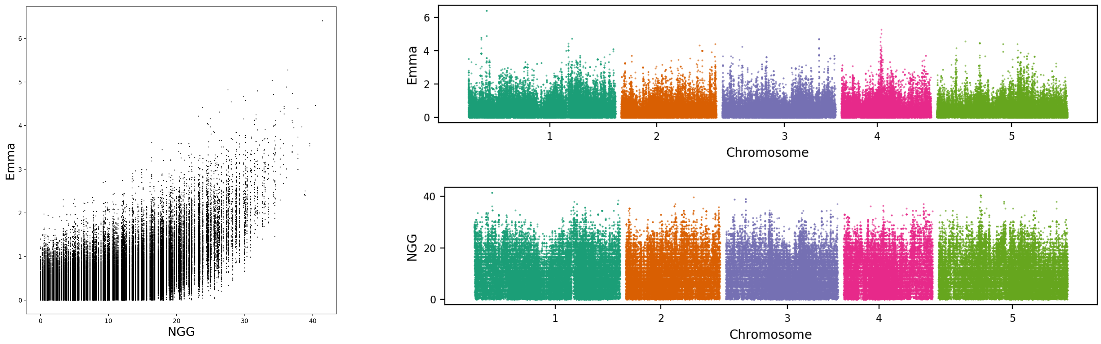

17
bacterial disease resistance
(avrRpm1)

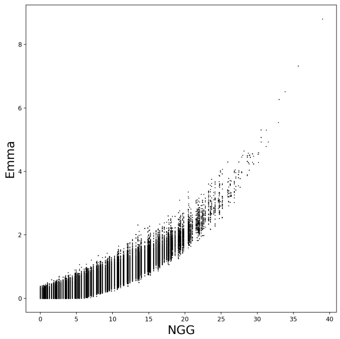


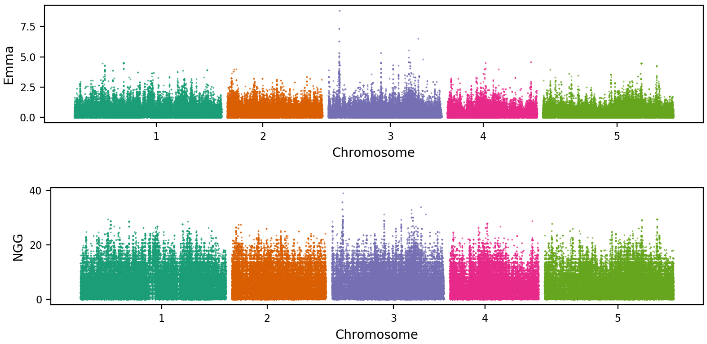


18
plant width
(Width 10)

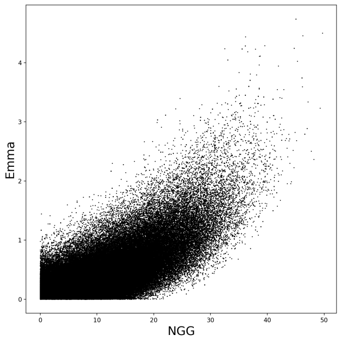


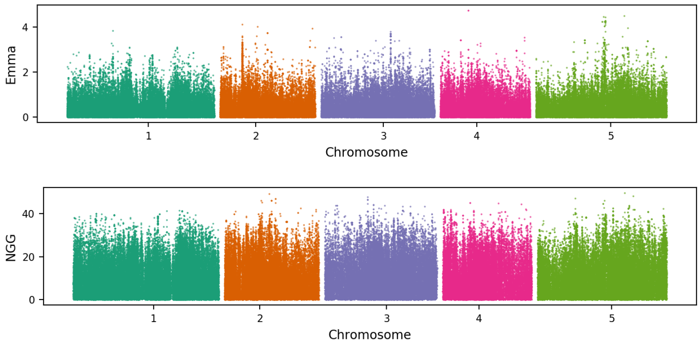


19
leaf chlorosis
(Chlorosis 22)

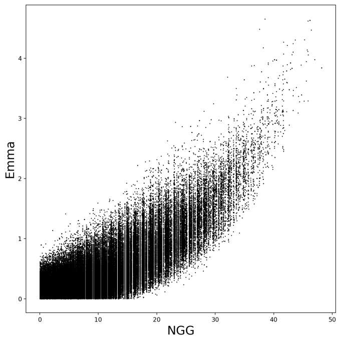


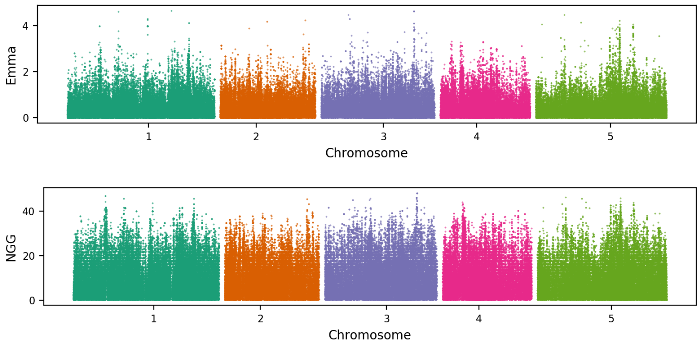


20
seed dormancy
(Storage 7 days)

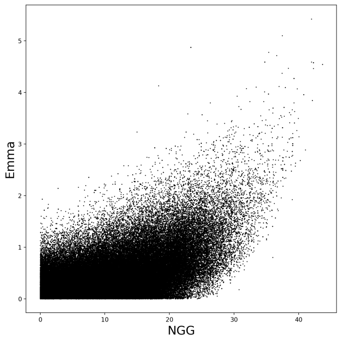


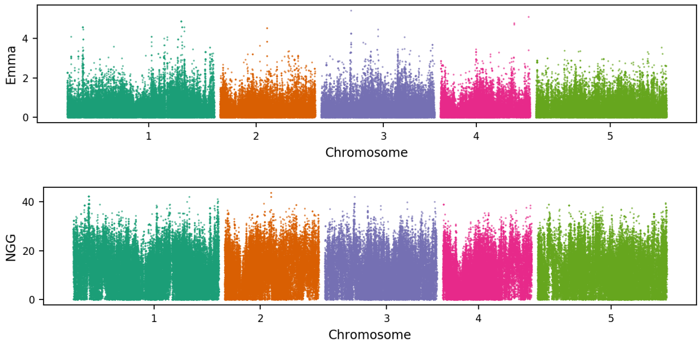

21
bacterial disease resistance
(As2 CFU2)

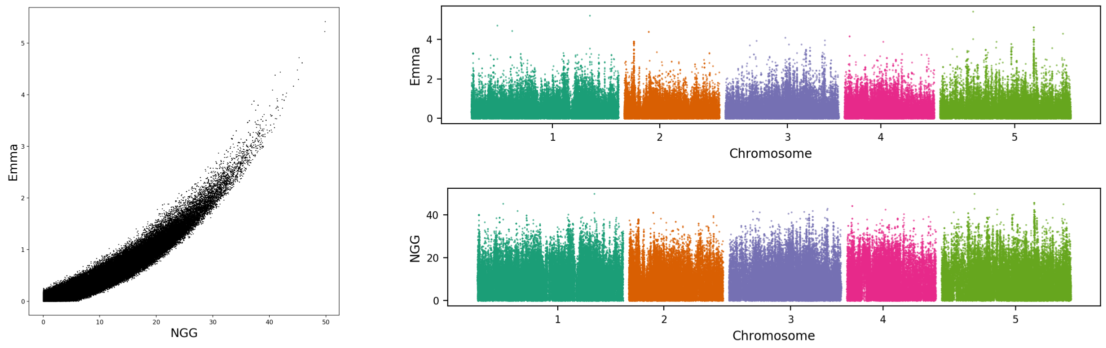


22
cobalt concentration
(Co59)

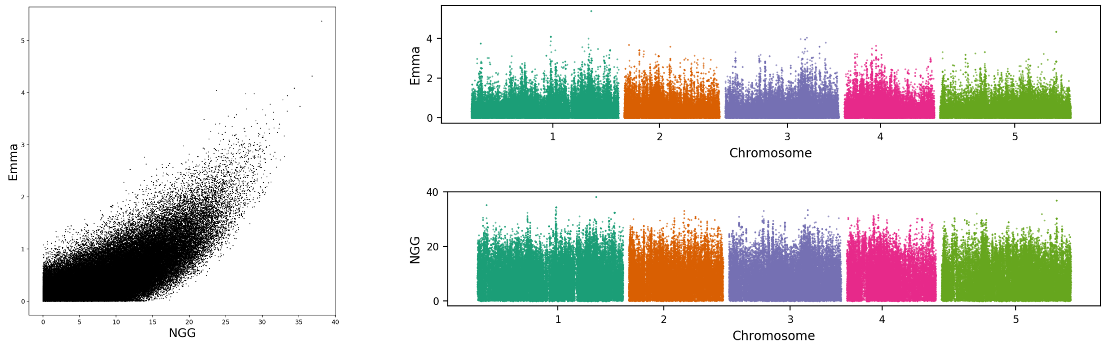


23
leaf necrosis
(FW)

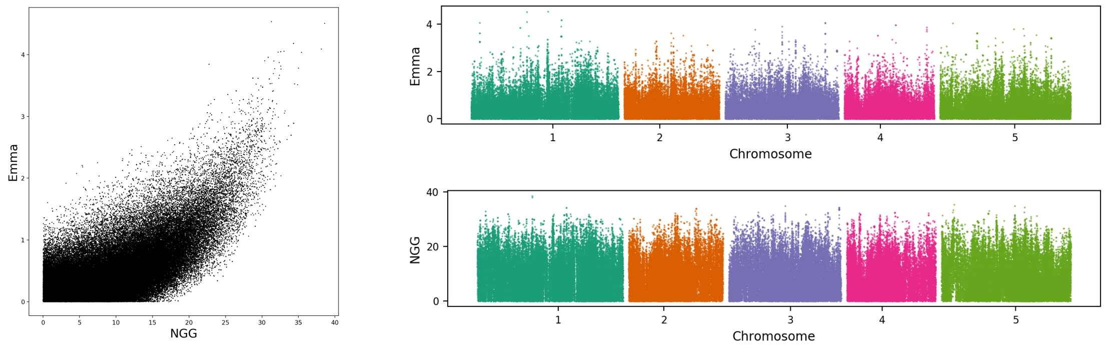


24
copper concentration
(Cu65)

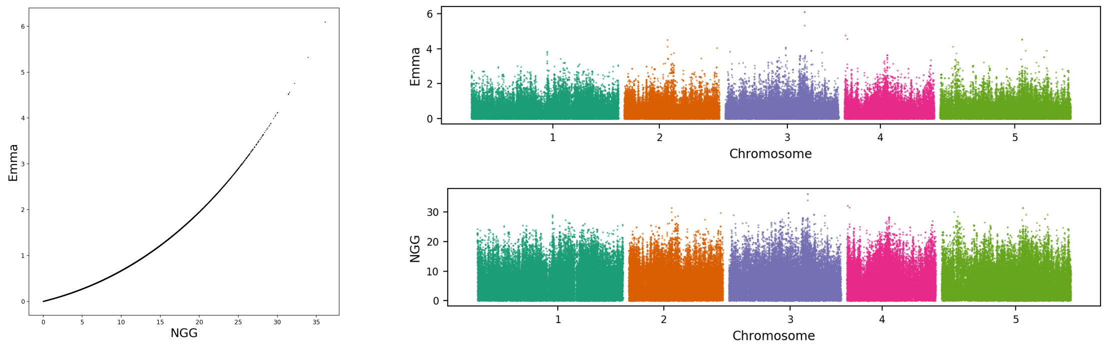

25
bacterial disease resistance
(Bacterial titer)

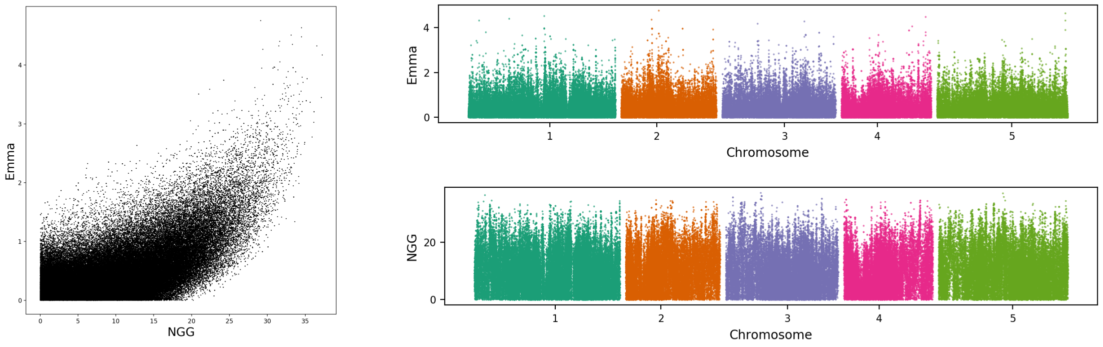


26
plant width
(Width 22)

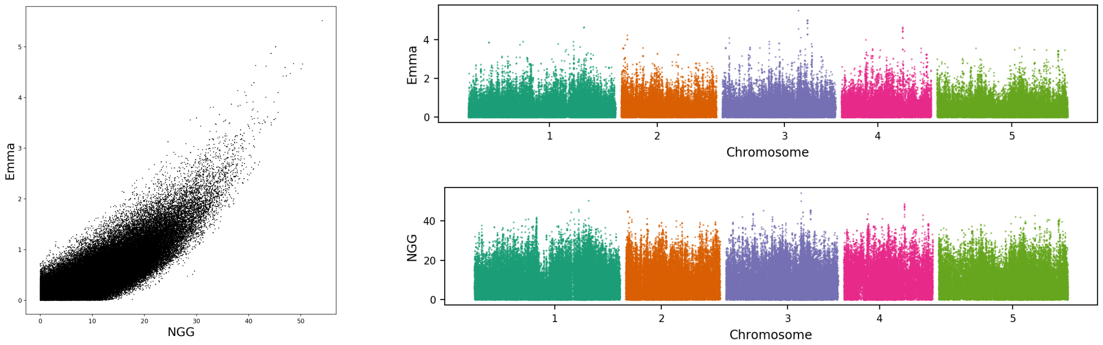


27
seed dormancy
(Storage 56 days)

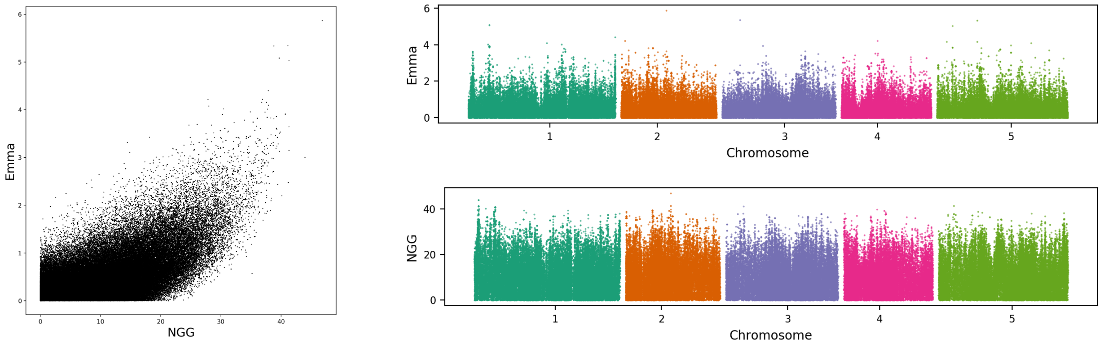


28
leaf chlorosis
(YEL)

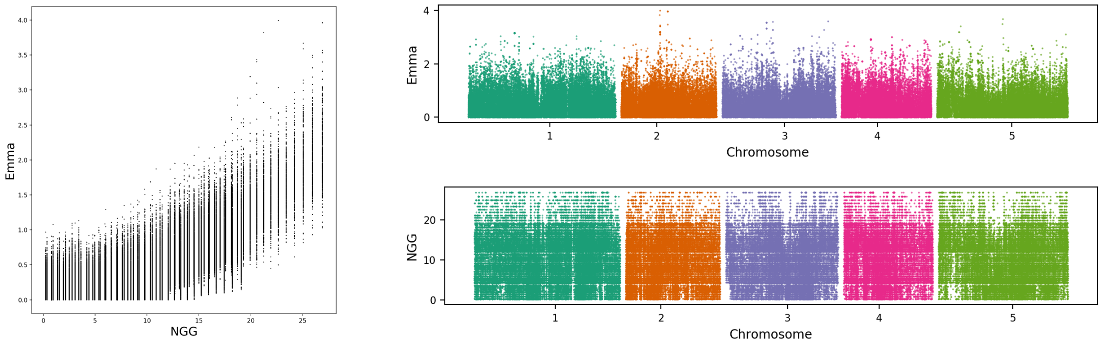

29
flowering time trait
(FLC)

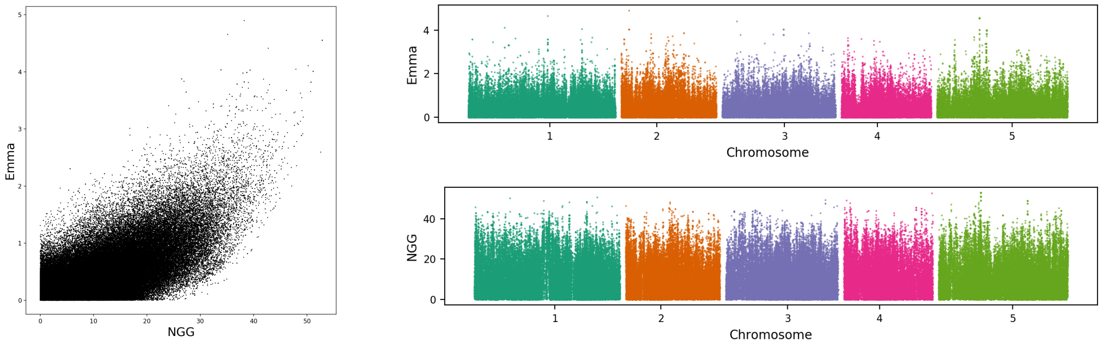


30
days to flowering
trait
(FT16)

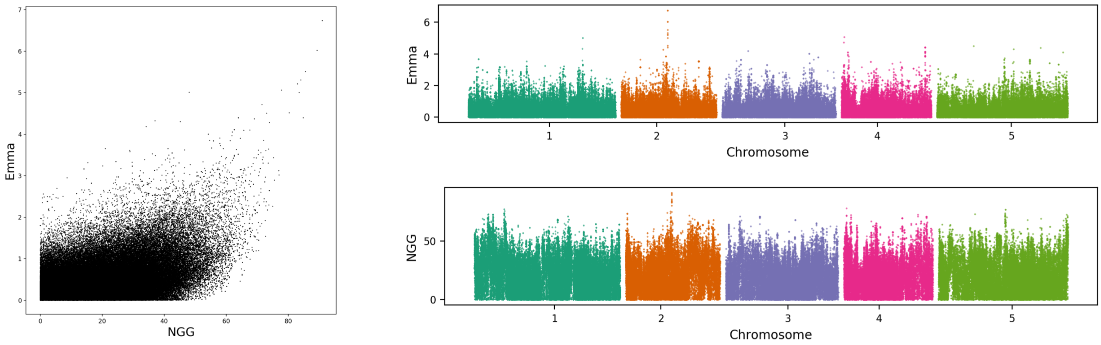


31
days to flowering
trait
(FT10)

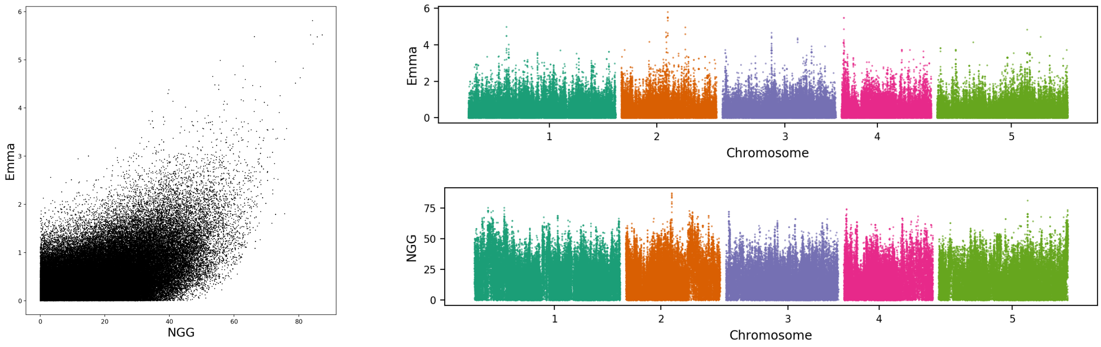


32
reproductive growth
time
(FT Duration GH)

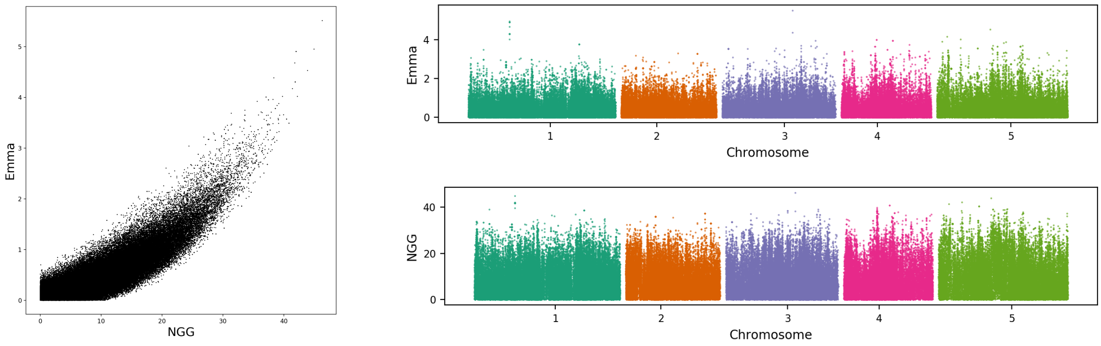

33
selenium
concentration
(Se82)

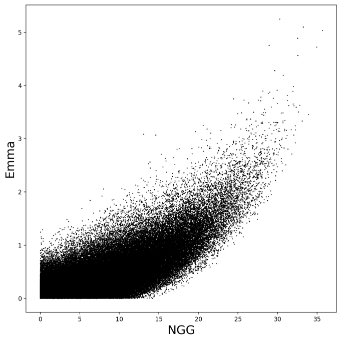


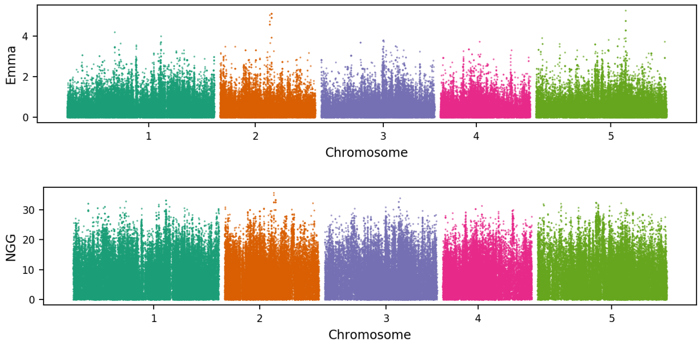


34
days to flowering
trait
(LDV)

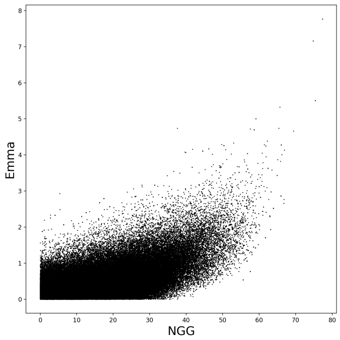


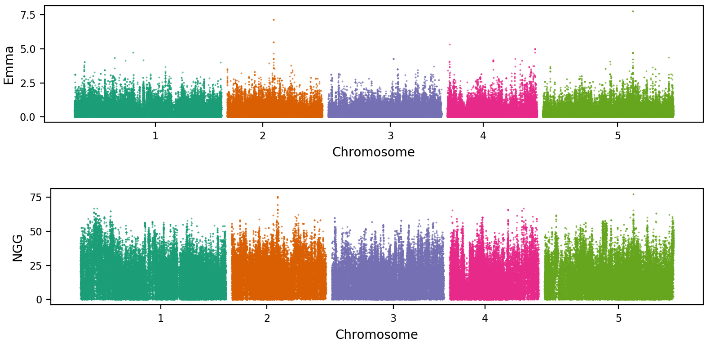


35
protist disease
resistance
(Noco2)

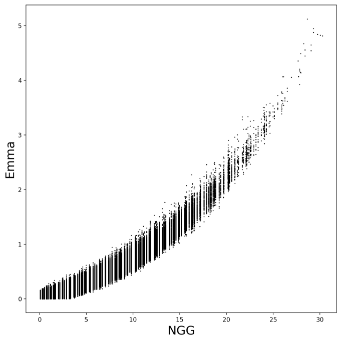


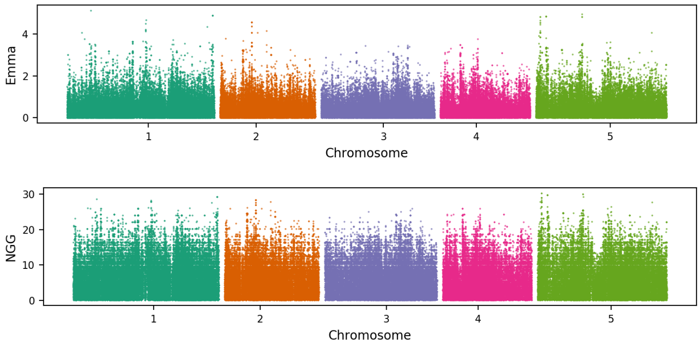


36
leaf number
(8W GH LN)

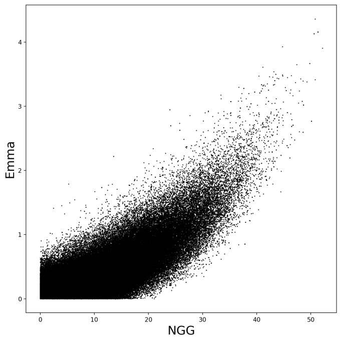


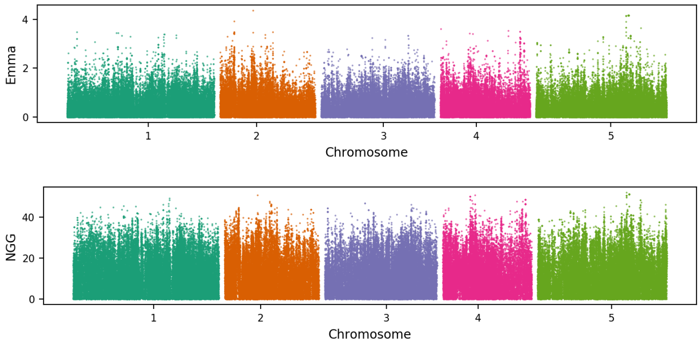

37
days to flowering
trait
(0W)

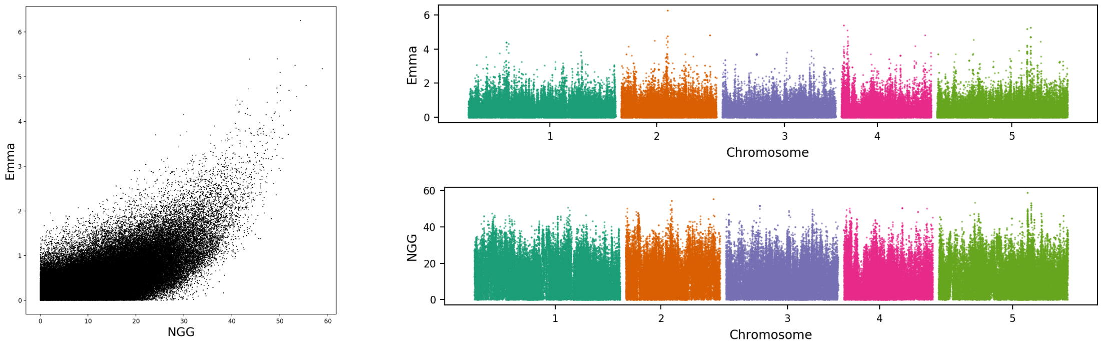


38
reproductive growth
time
(MT GH)

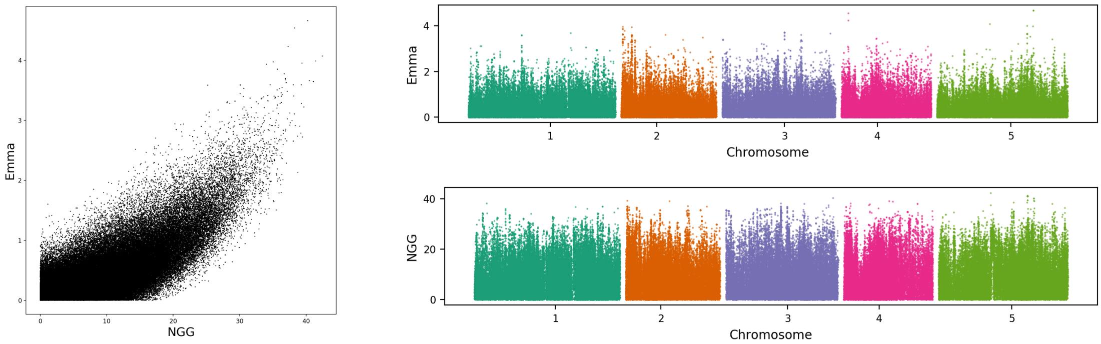


39
relative growth rate
(After Vern Growth)

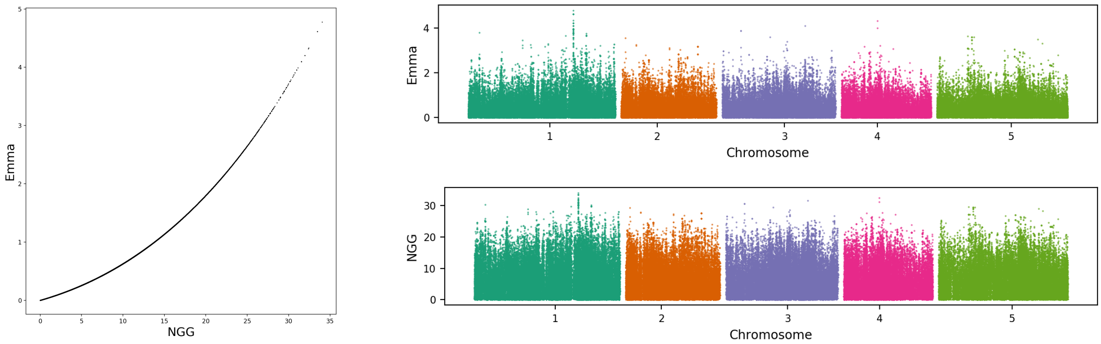


40
aphid resistance
(Aphid number)

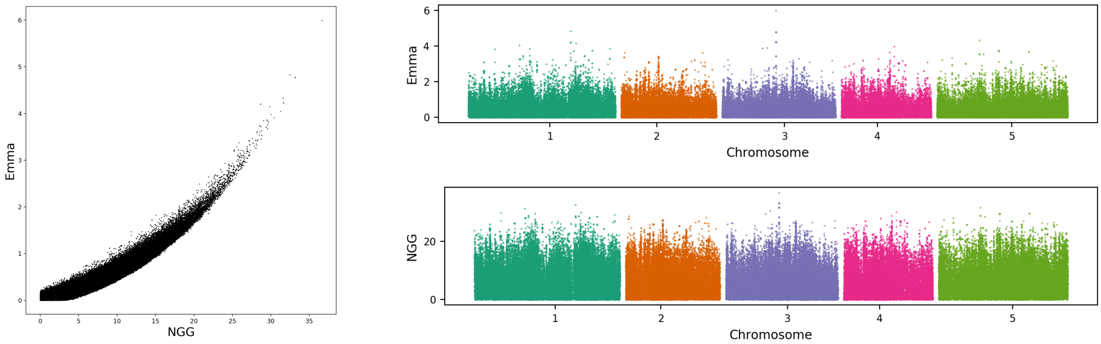

41
leaf number
(LN22)

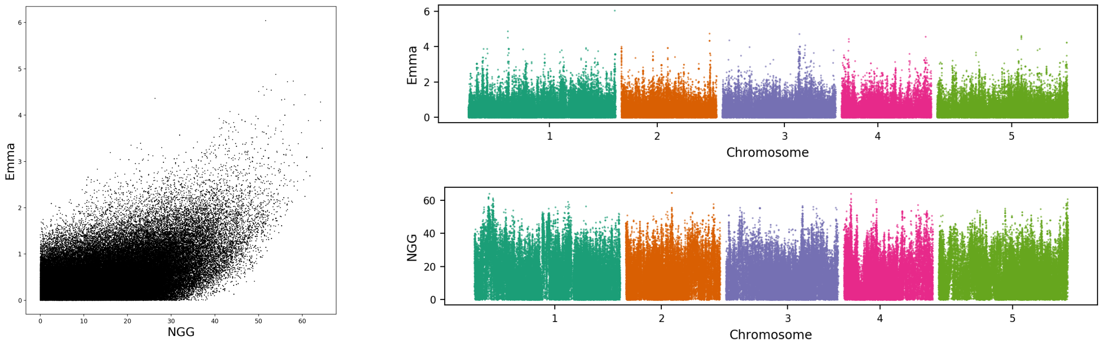


42
bacterial disease
resistance
(Bs CFU2)

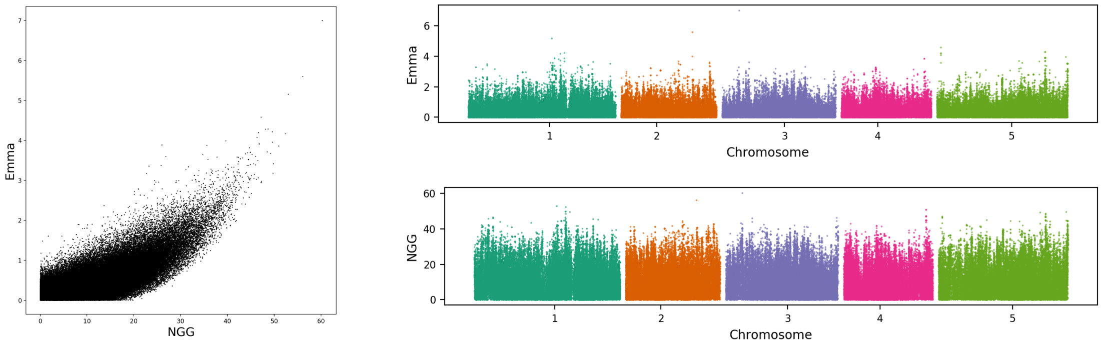


43
bacterial disease
resistance
(avrRpt2)

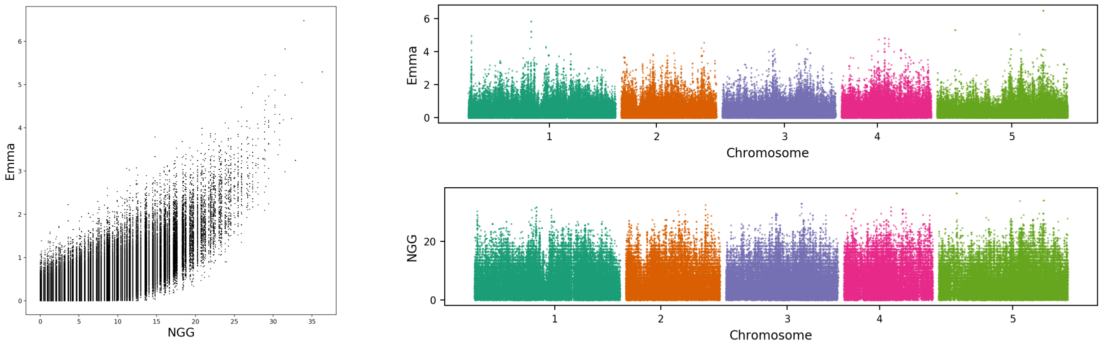


44
seedling hypocotyl
length
(Hypocotyl length)

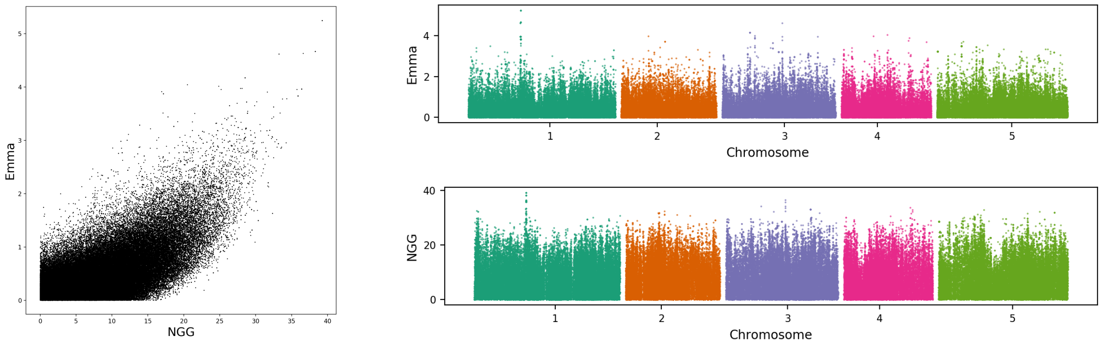

45
days to germinate
(Germ22)

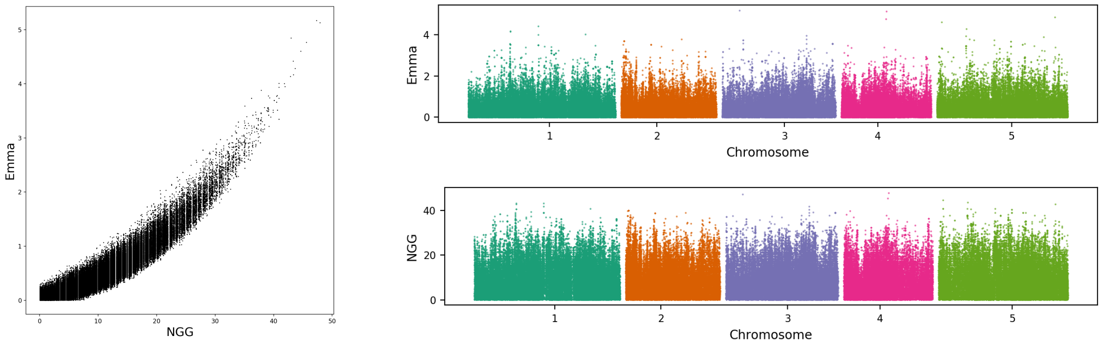


46
rolled leaf
(Leaf roll 22)

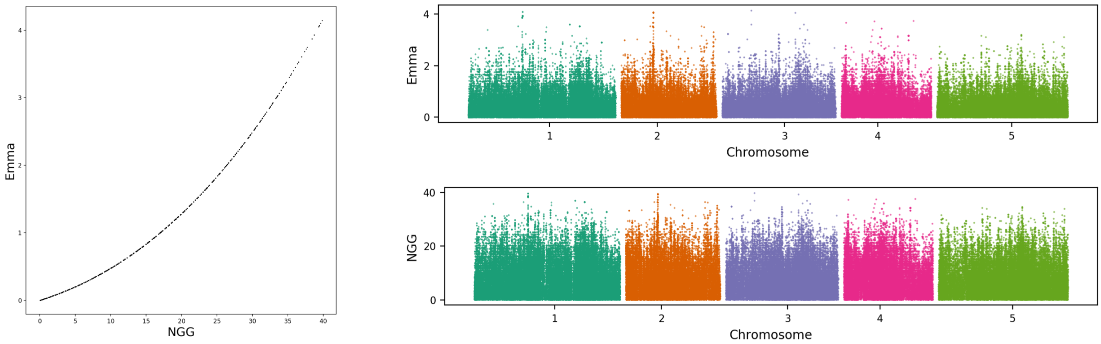


47
days to flowering
trait
(SD)

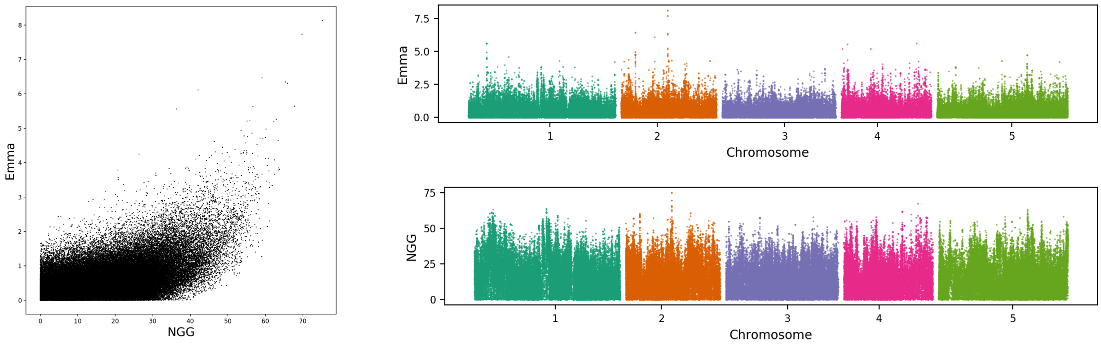


49
days to flowering
trait
(FT GH)

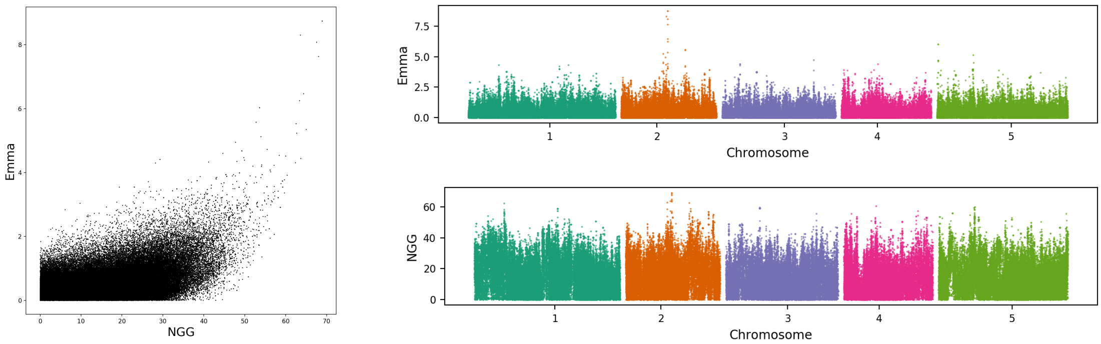

50
DSDS50
(DSDS50)

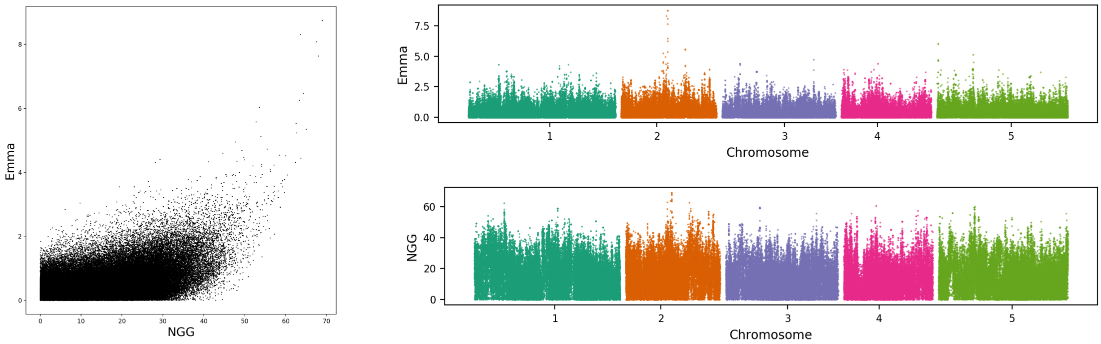


51
calcium
concentration
(Ca43)

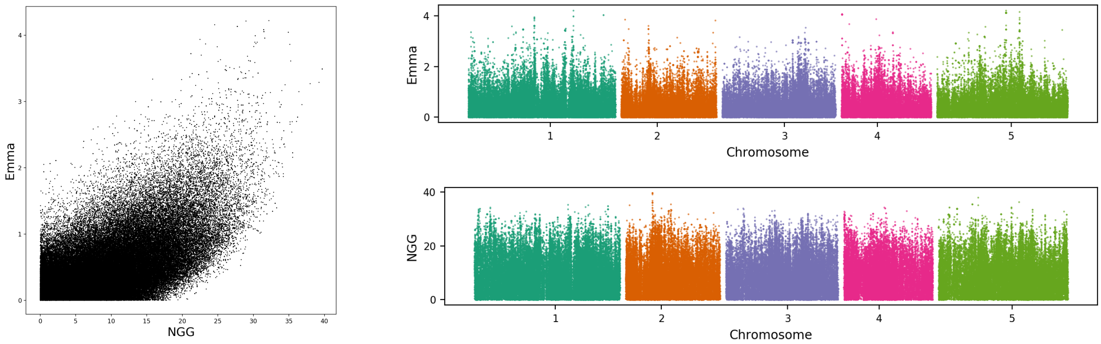


52
reproductive growth
time
(LC Duration GH)

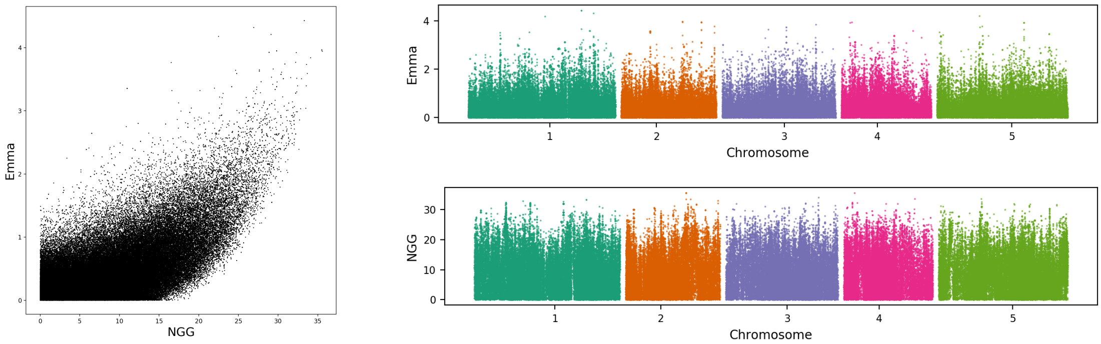


53
days to flowering
trait
(0W GH FT)

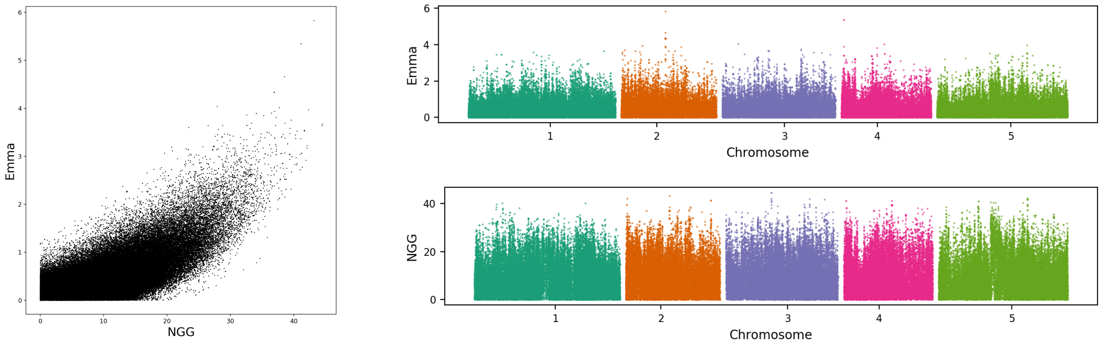

54
boron concentration
(B11)

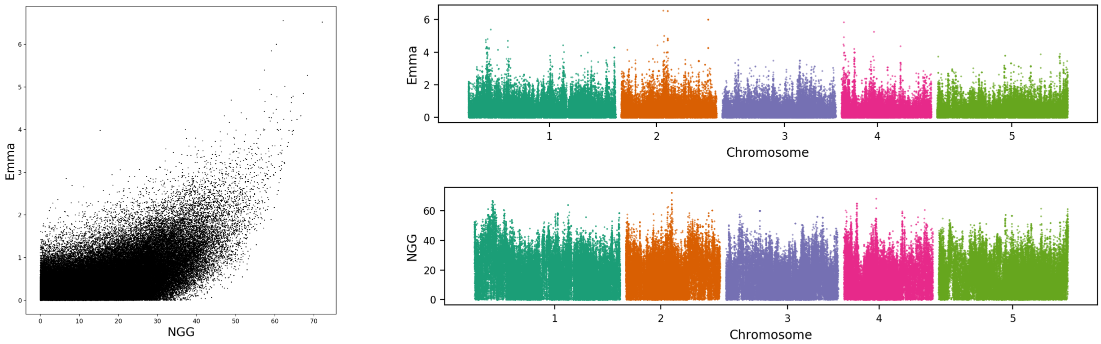


55
leaf chlorosis
(Chlorosis 10)

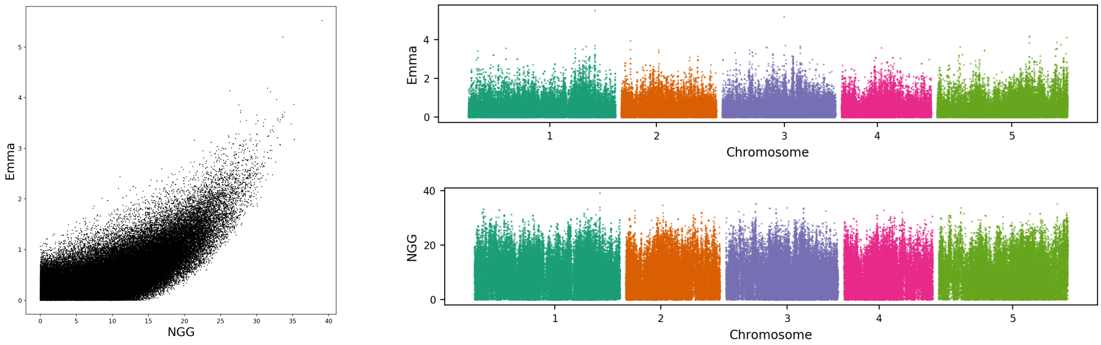


56
reproductive growth
time
(RP GH)

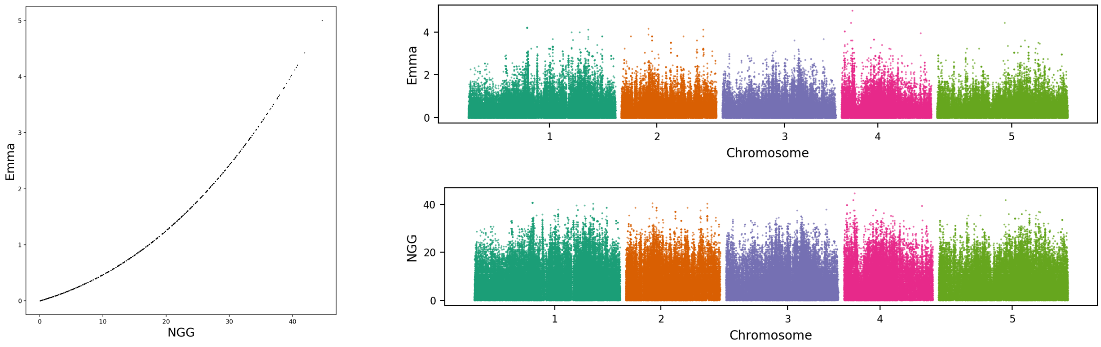


57
leaf chlorosis
(Chlorosis 16)

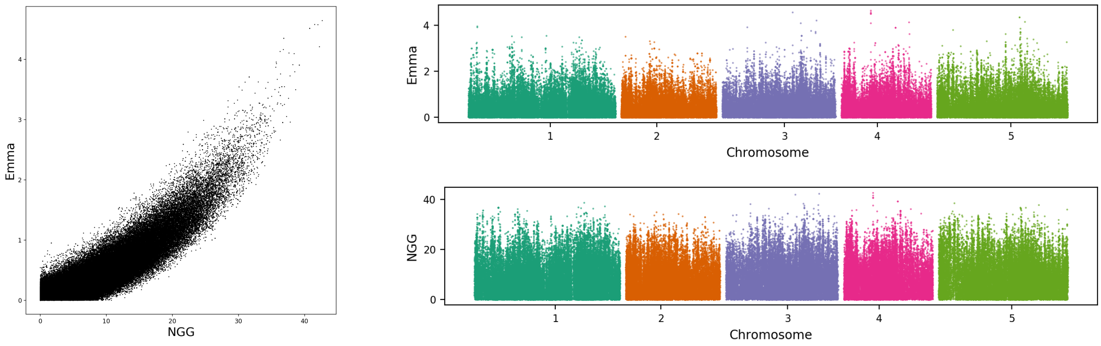

58
reproductive growth time
(LFS GH)

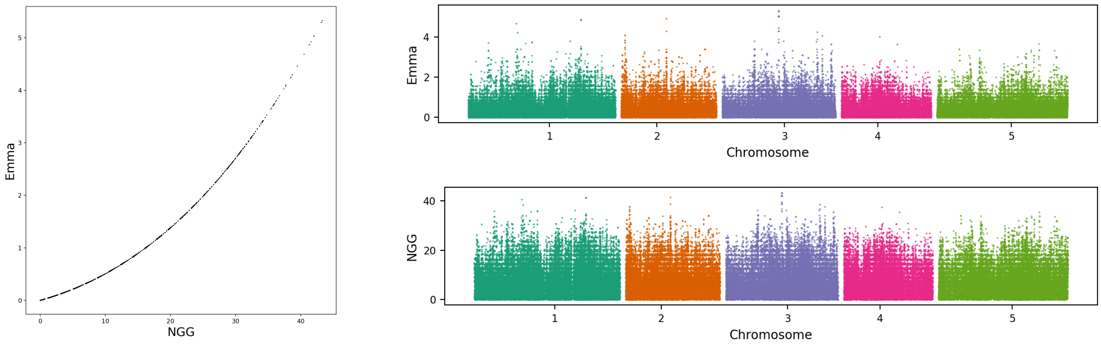


59
days to germinate
(Germ10)

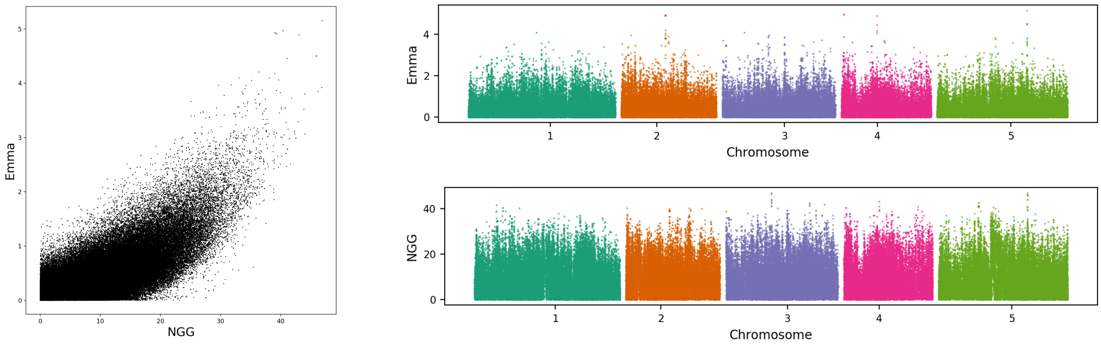


60
days to germinate
(Germ16)

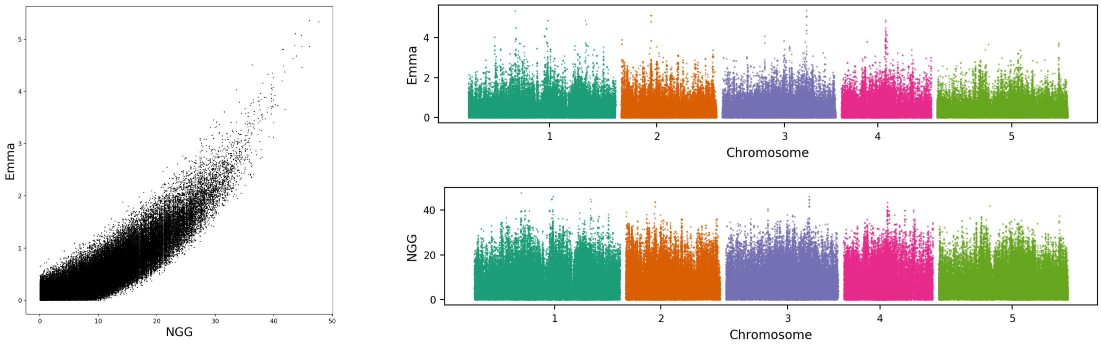


61
anthocyanin content
(Anthocyanin 16)

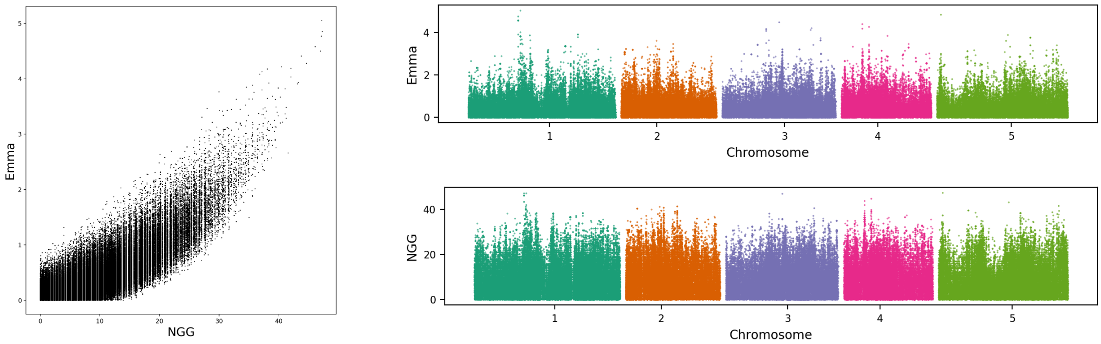

62
anthocyanin content
(Anthocyanin 10)

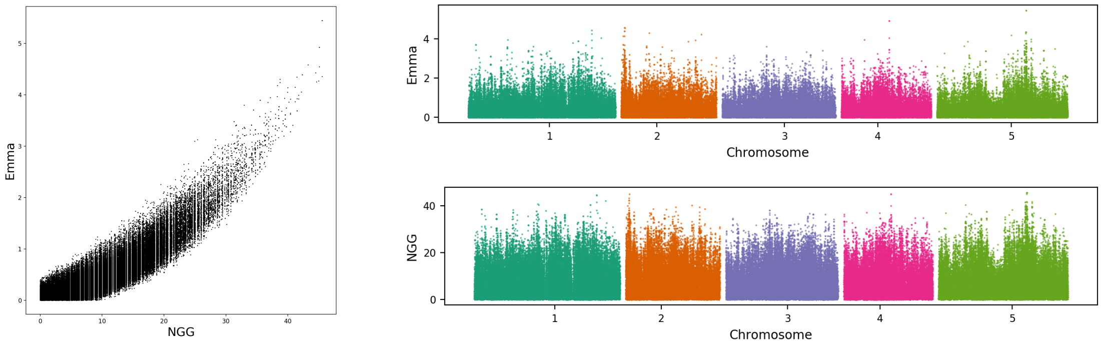


63
bacterial disease
resistance
(At1 CFU2)

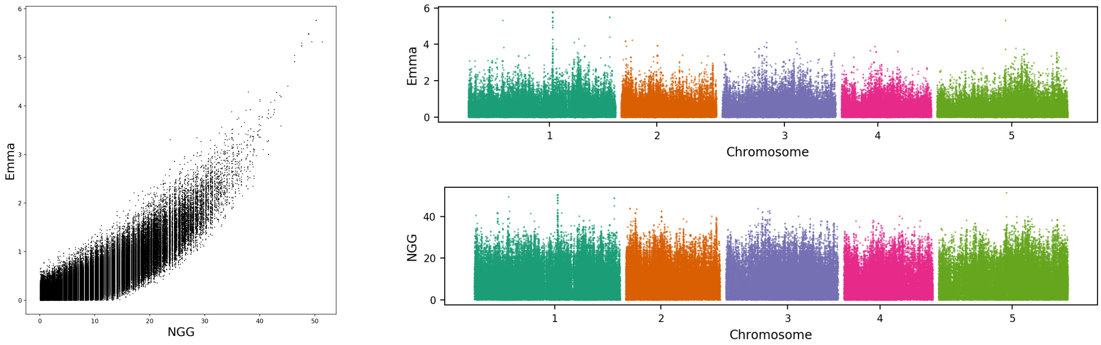


64
nickel concentration
(Ni60)

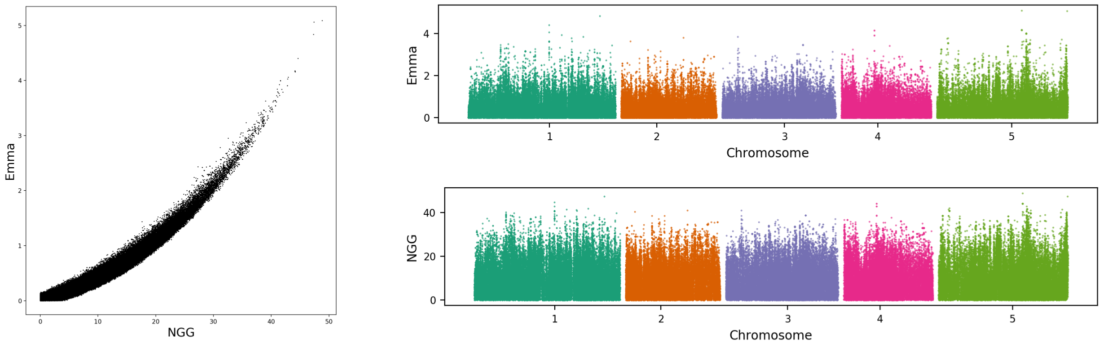


65
phosphorus
concentration
(P31)

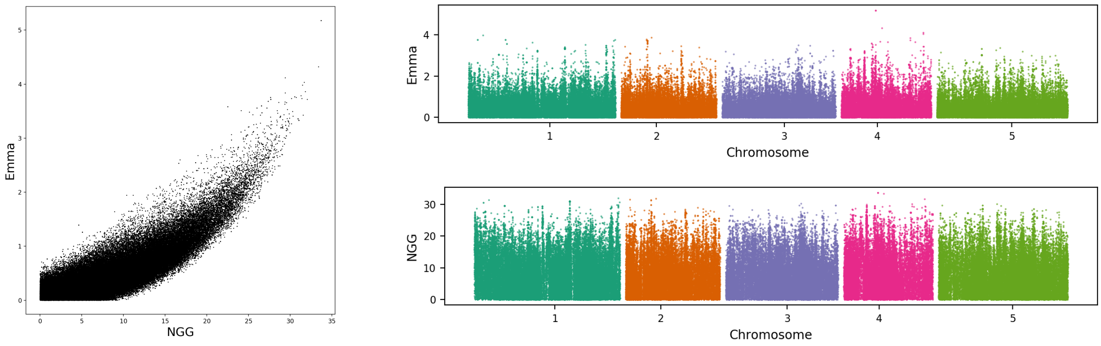

66
protist disease resistance
(Emwa1)

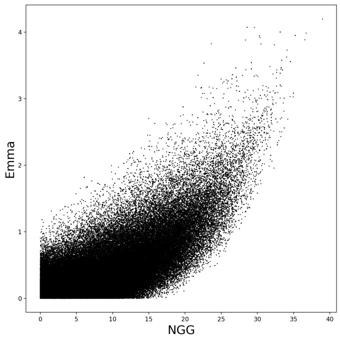


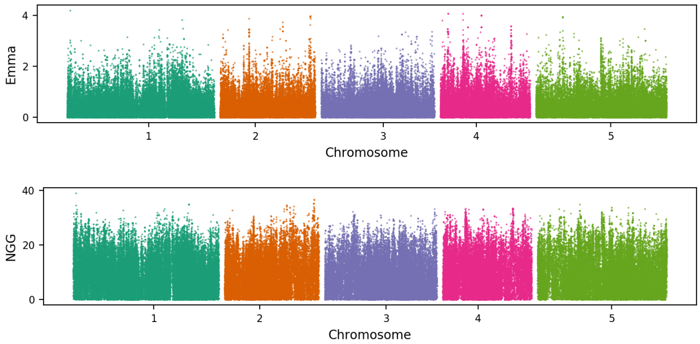


67
arsenic concentration
(As75)

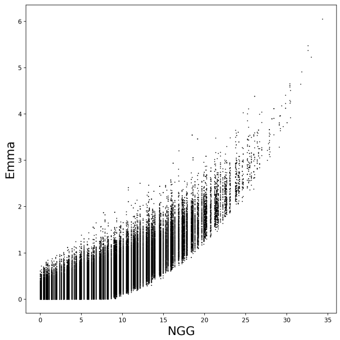


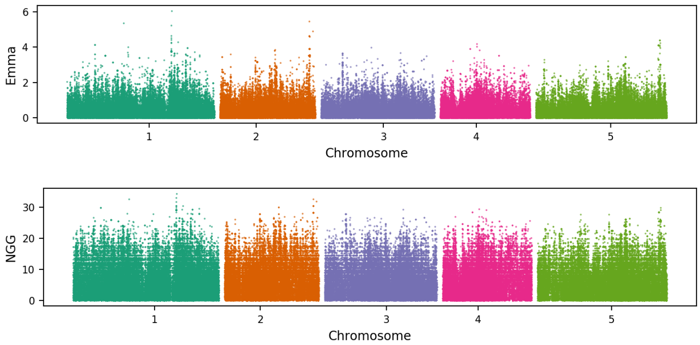


68
germinability in dark
(Germ in dark)

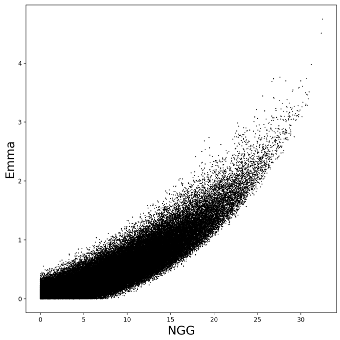


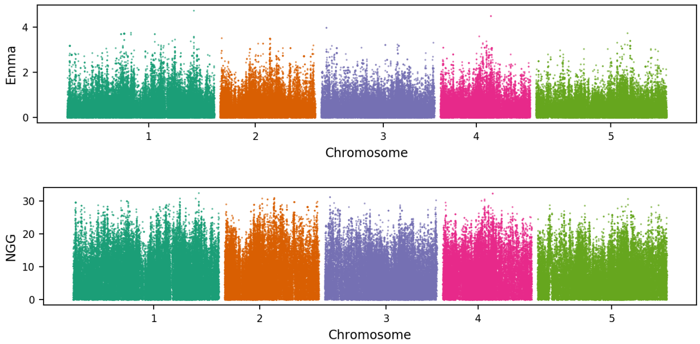


69
flowering time trait
(FRI)

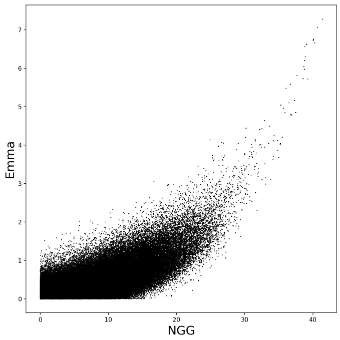


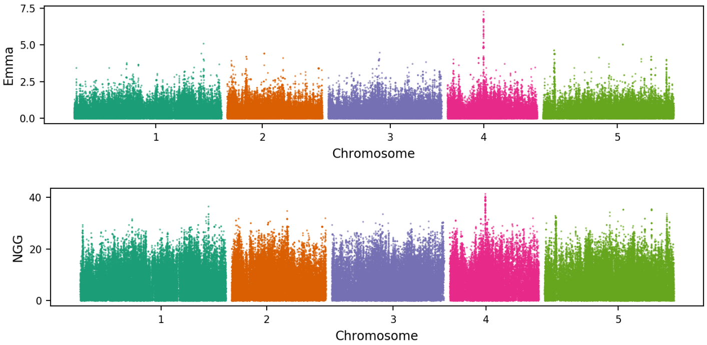

70
bacterial disease resistance
(As CFU2)

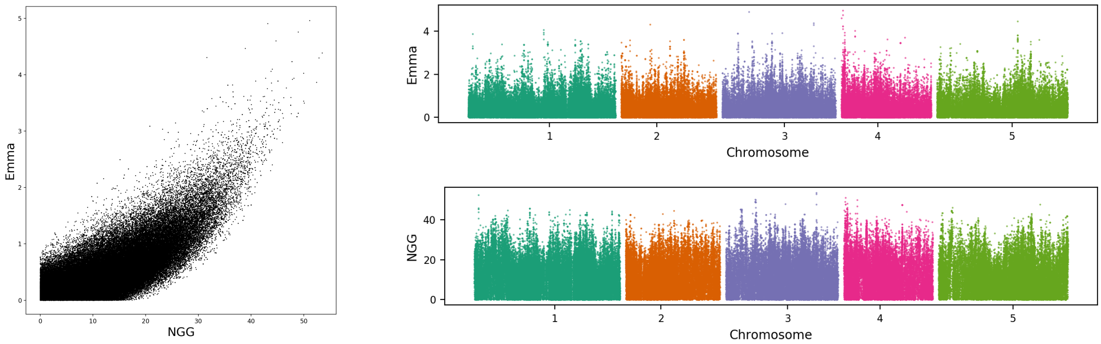


71
leaf trichome density
(Trichome avg C)

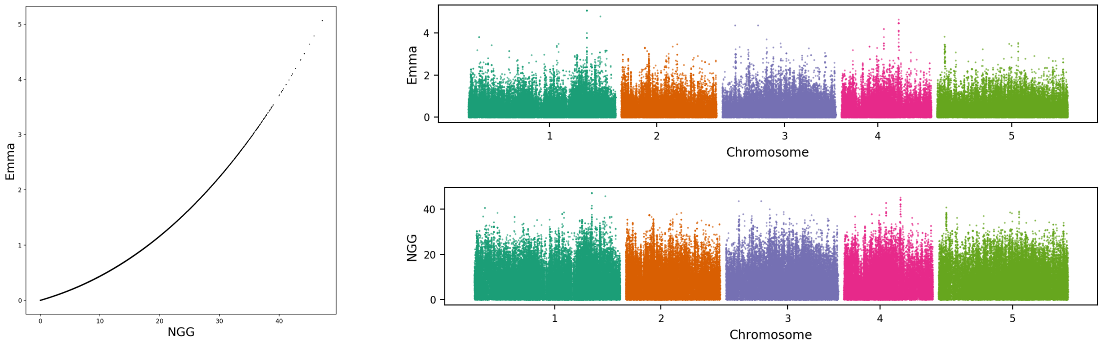


72
relative growth rate
(Vern Growth)

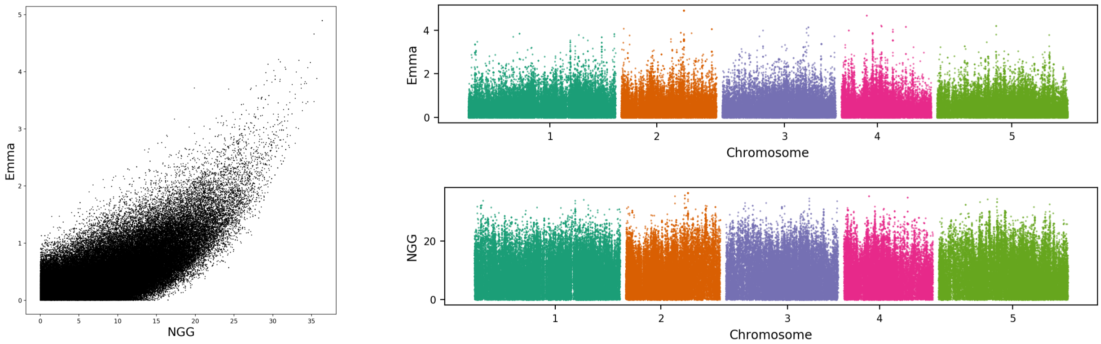


73
molybdenum concentration
(Mo98)

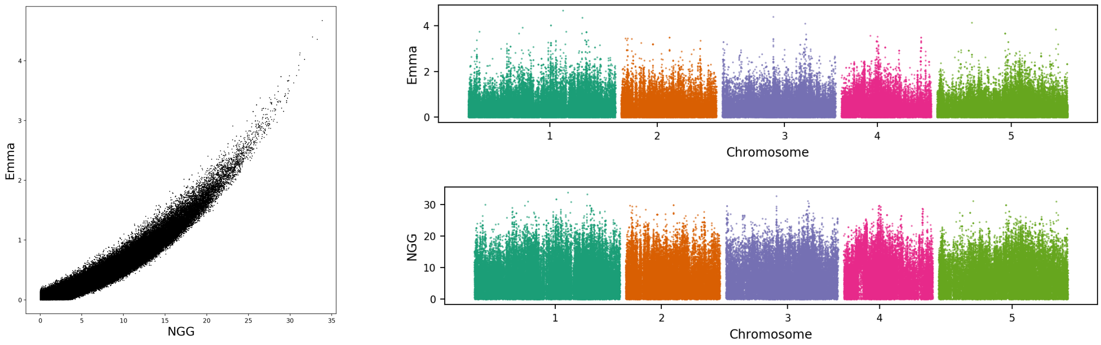

74
protist disease resistance
(Hiks1)

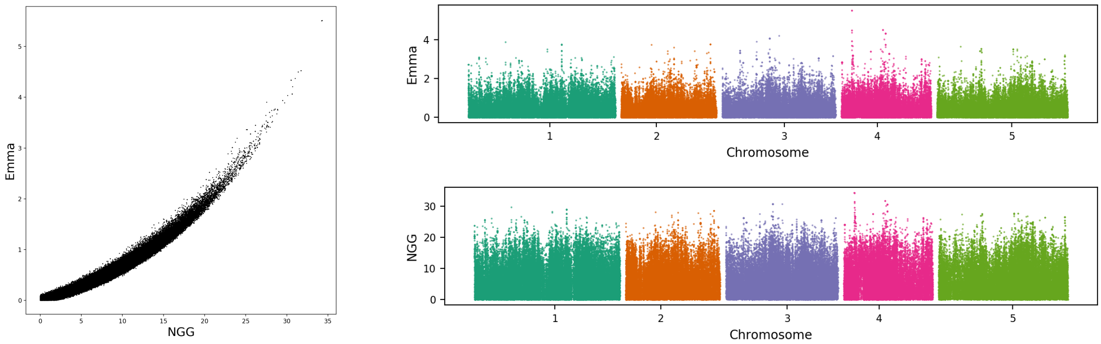


75
anthocyanin content
(Anthocyanin 22)

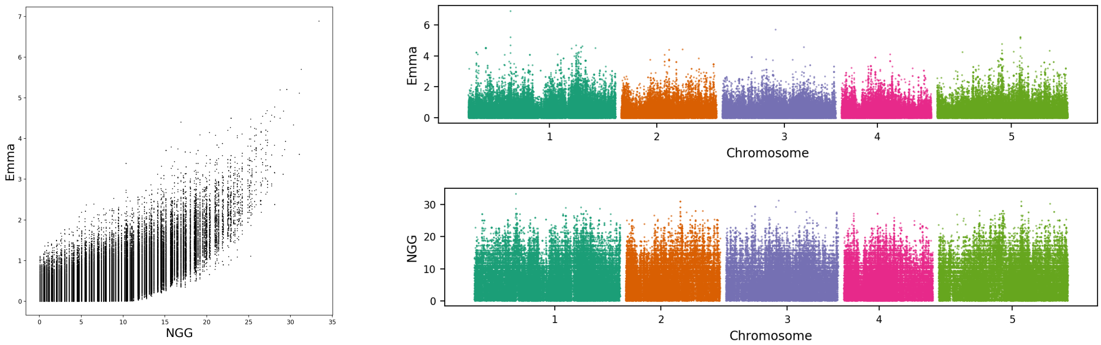


76
zinc concentration
(Zn66)

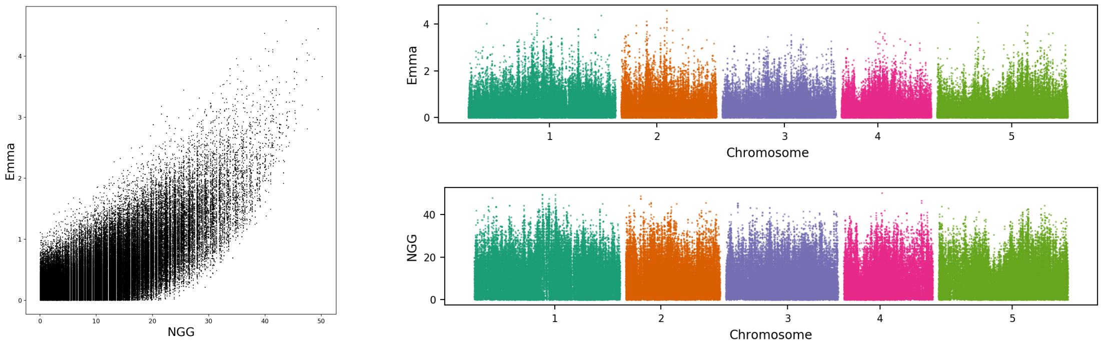


77
leaf trichome density
(Trichome avg JA)

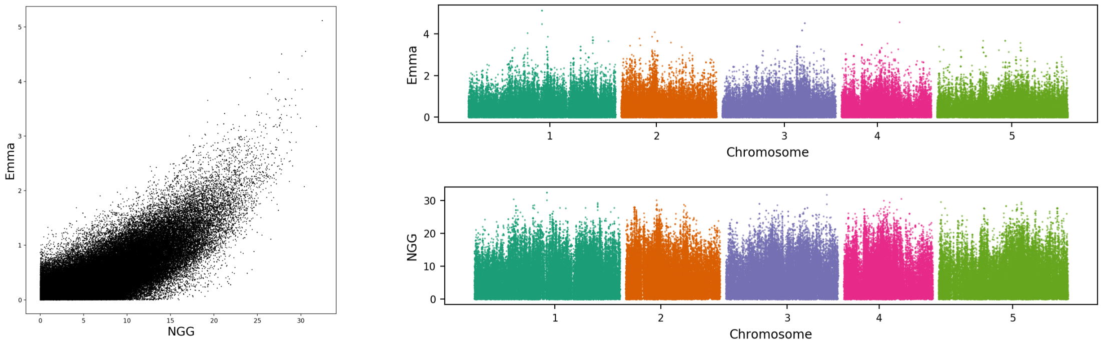

78
leaf necrosis
(LES)

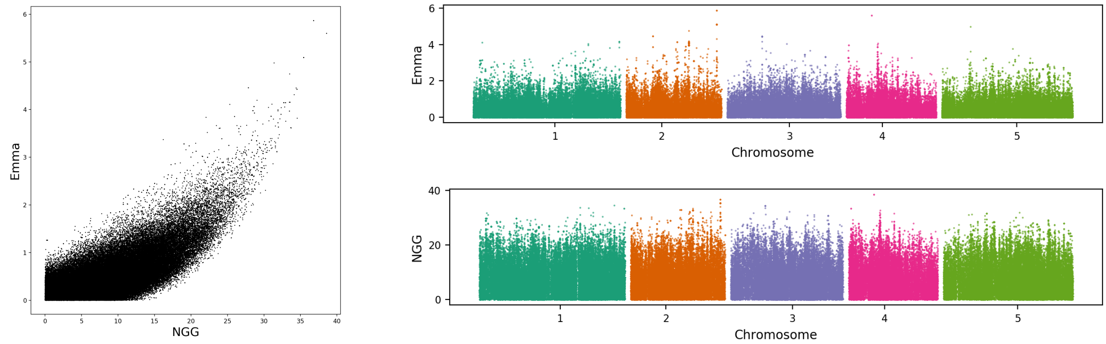


79
fruit length
(Silique 16)

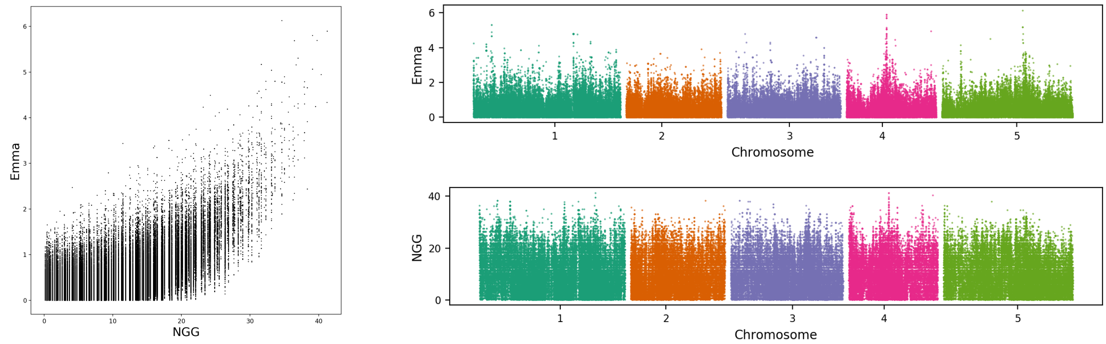


80
protist disease
resistance
(Emoy)

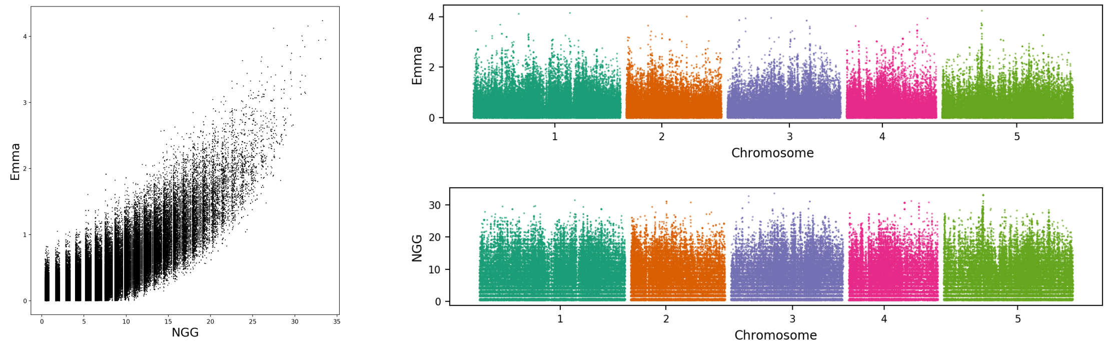


81
potassium
concentration
(K39)

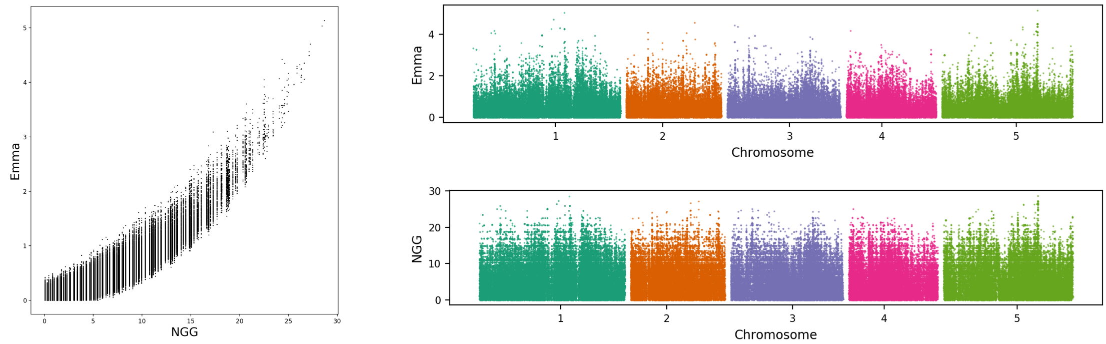

82
leaf number
(0W GH LN)

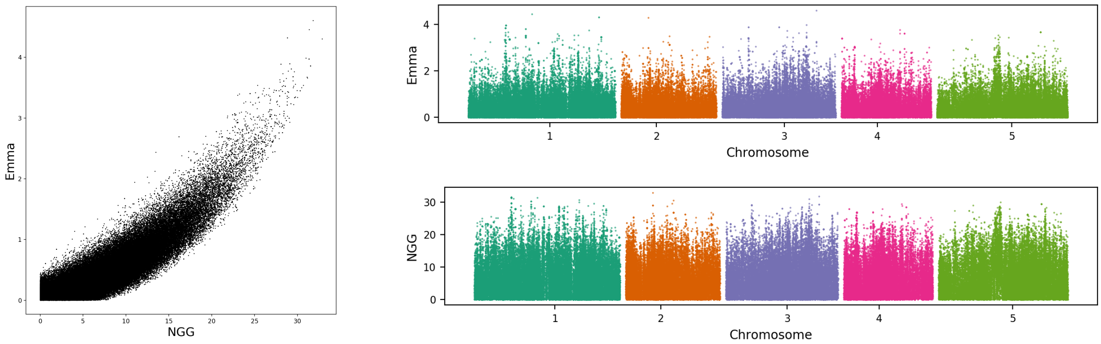


83
bacterial disease
resistance
(At2)

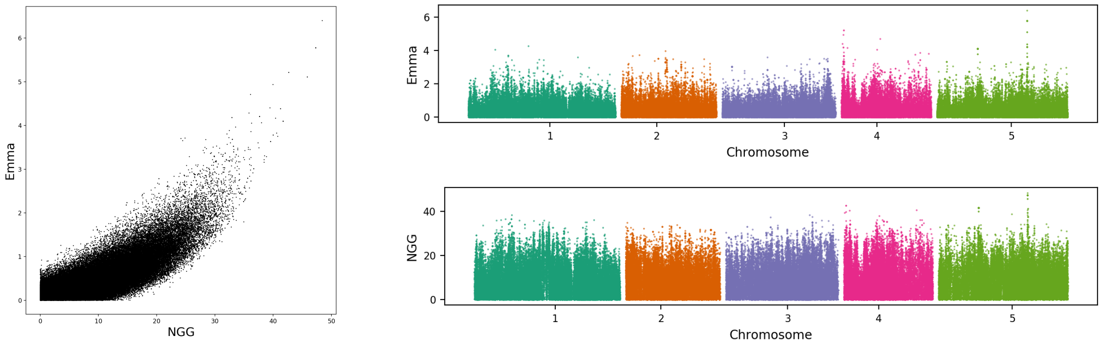


84
bacterial disease
resistance
(At1)

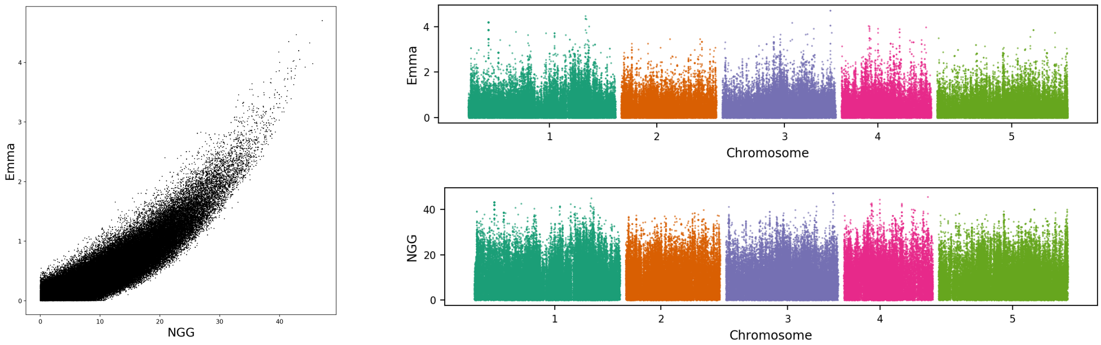


85
leaf number
(LN10)

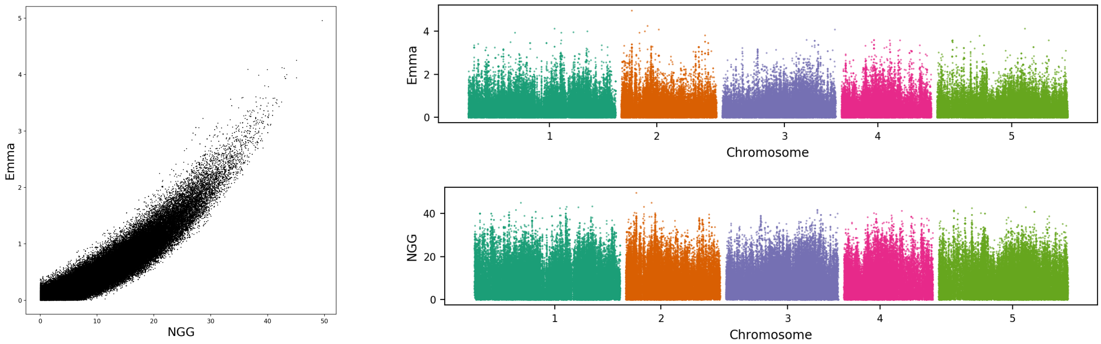

86
days to flowering
trait
(FT Field)

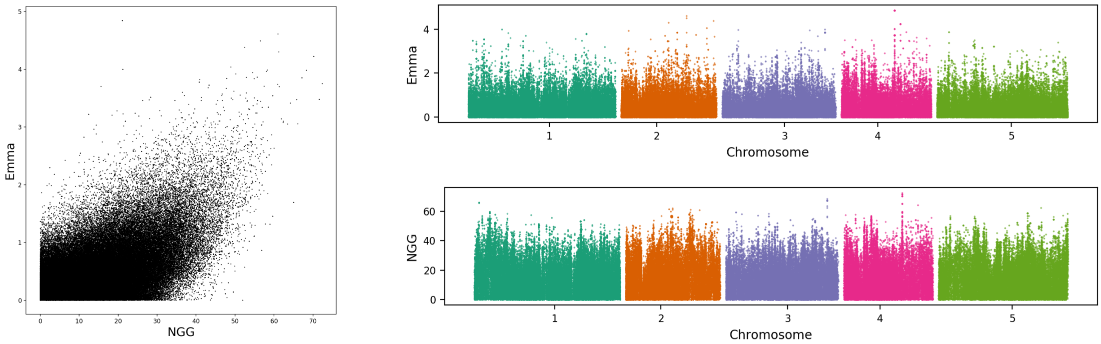


87
leaf number
(LN16)

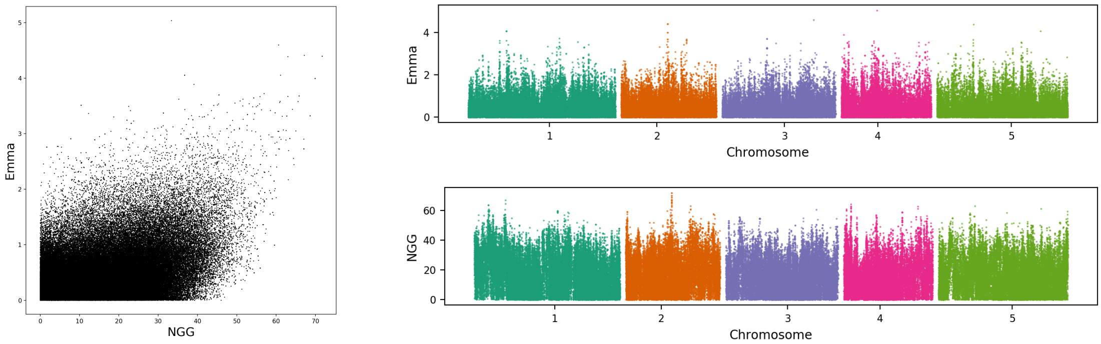


89
days to flowering
trait
(LD)

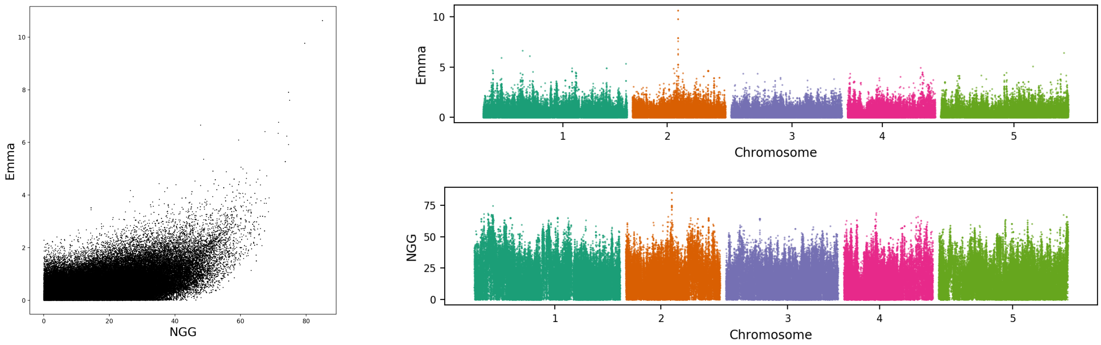


90
relative growth rate
(Seedling Growth)

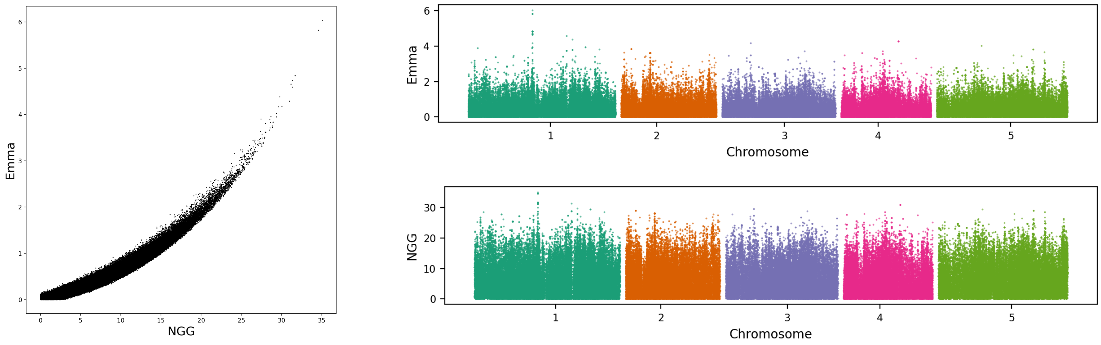

91
sulfur concentration
(S34)

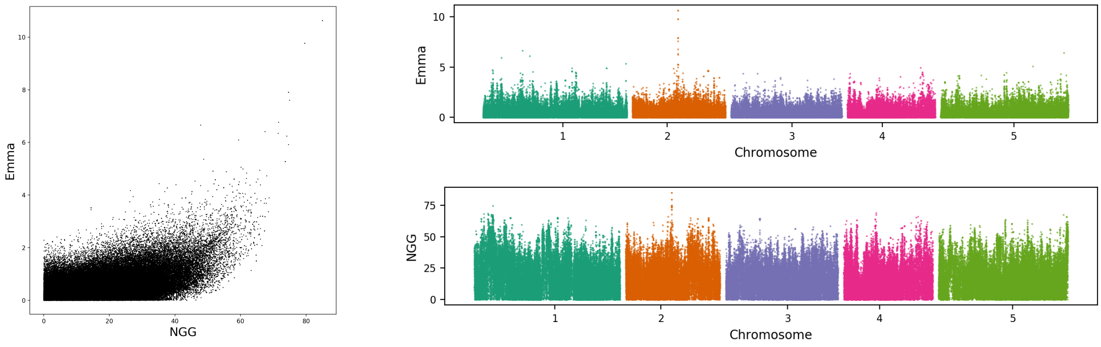


92
leaf margin serrated
(Leaf serr 22)

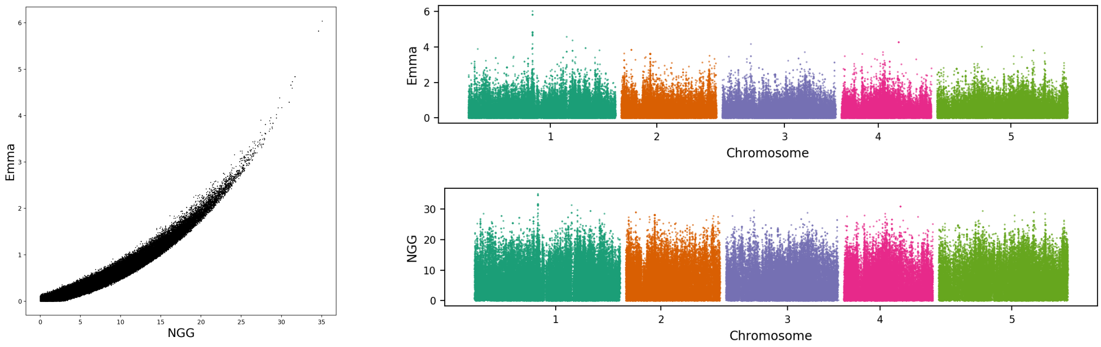


93
leaf necrosis
(DW)

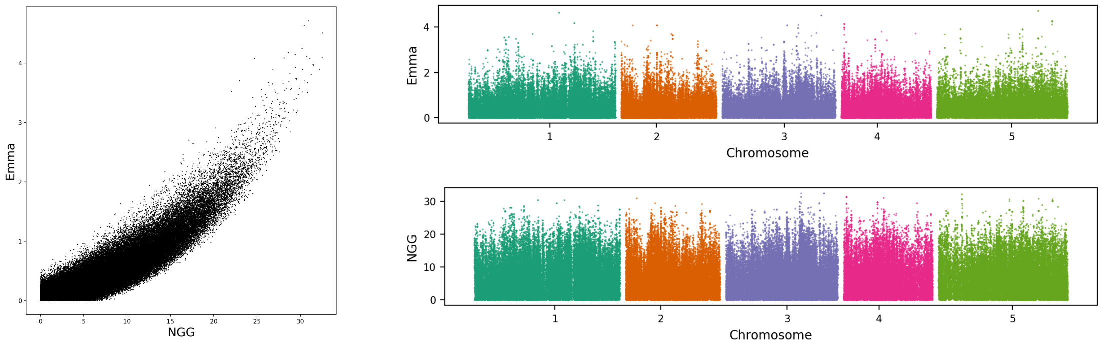


94
seed dormancy
(Seed Dormancy)

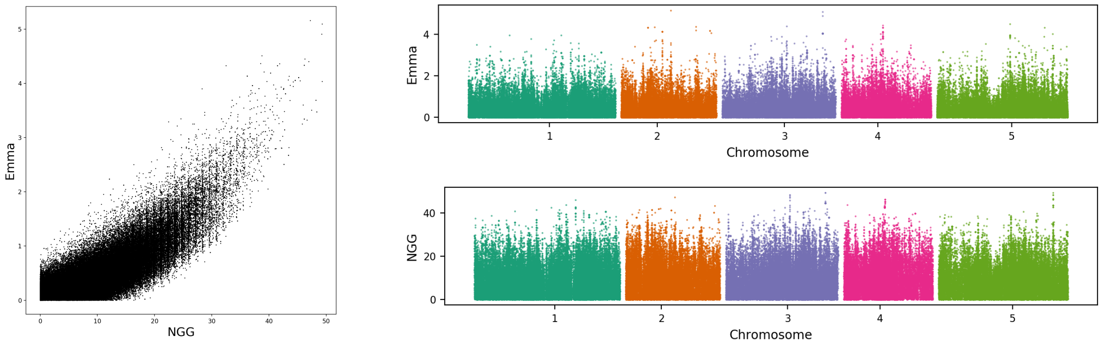

95
manganese
concentration
(Mn55)

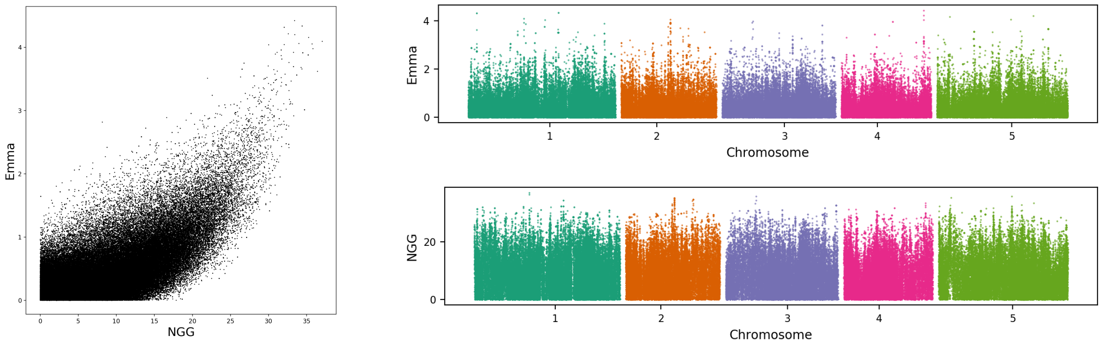


96
fruit length
(Silique 22)

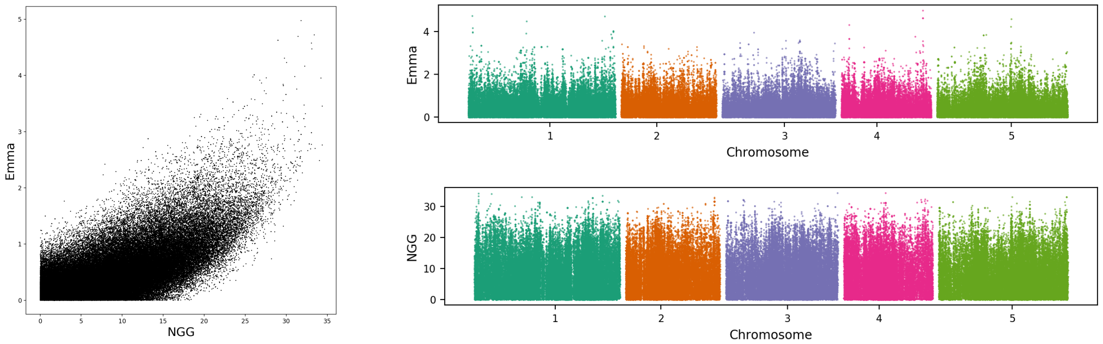


97
bacterial disease
resistance
(avrPphB)

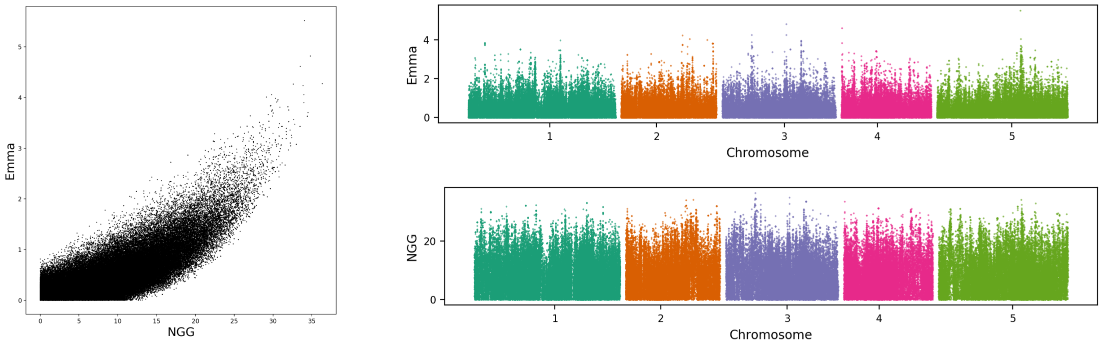


98
iron concentration
(Fe56)

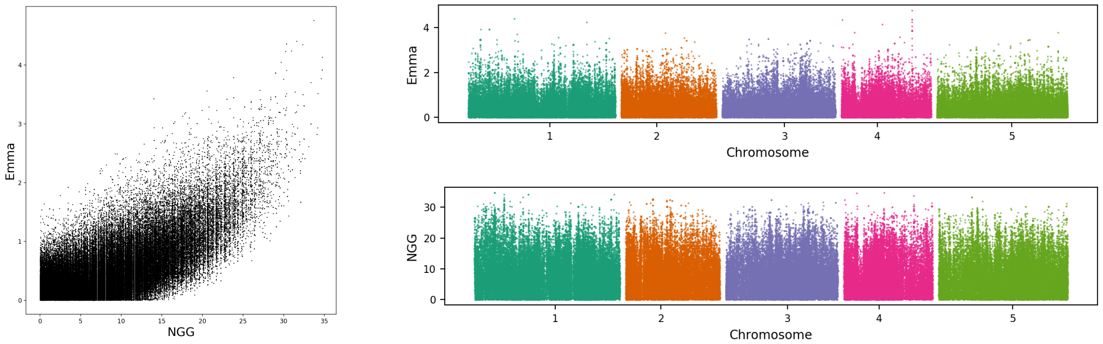

99
days to flowering
trait
(8W GH FT)

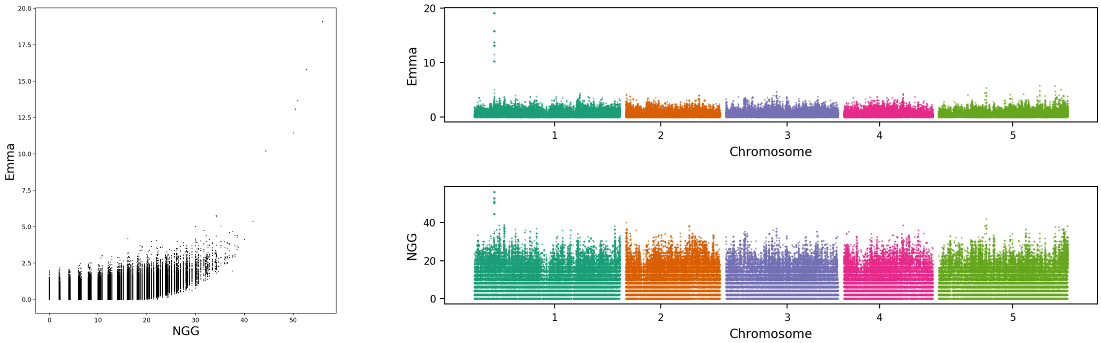


100
days to flowering
trait
(4W)

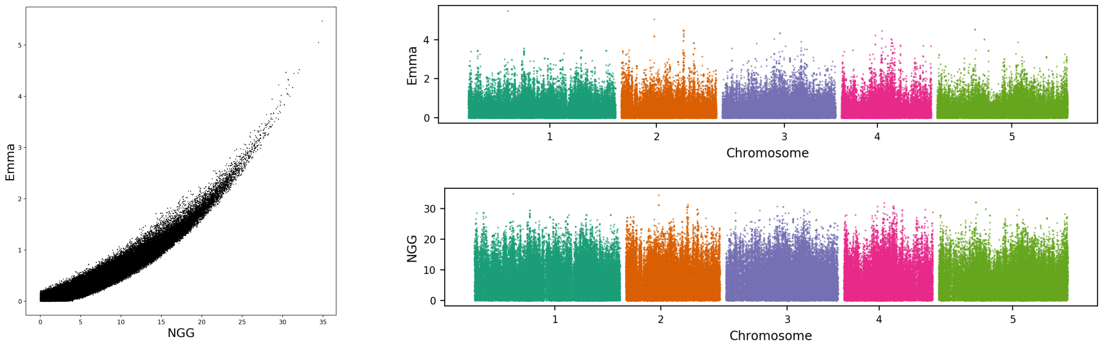


101
lithium
concentration
(Li7)

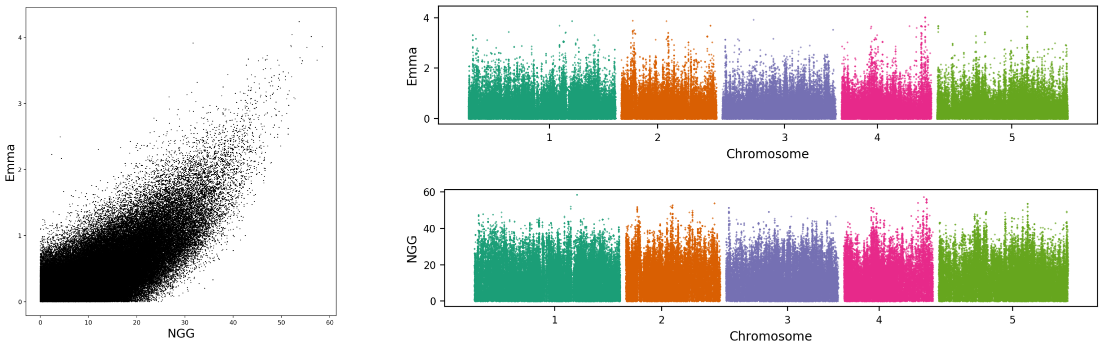


102
days to flowering
trait
(FT22)

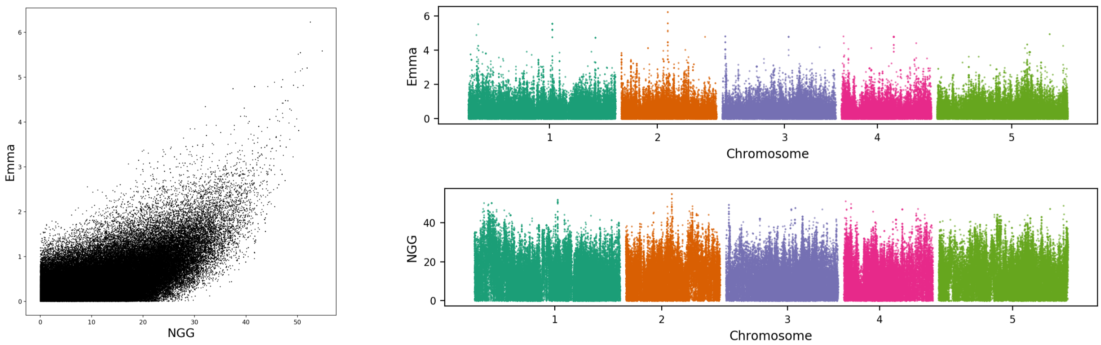

103
bacterial disease resistance
(As2)

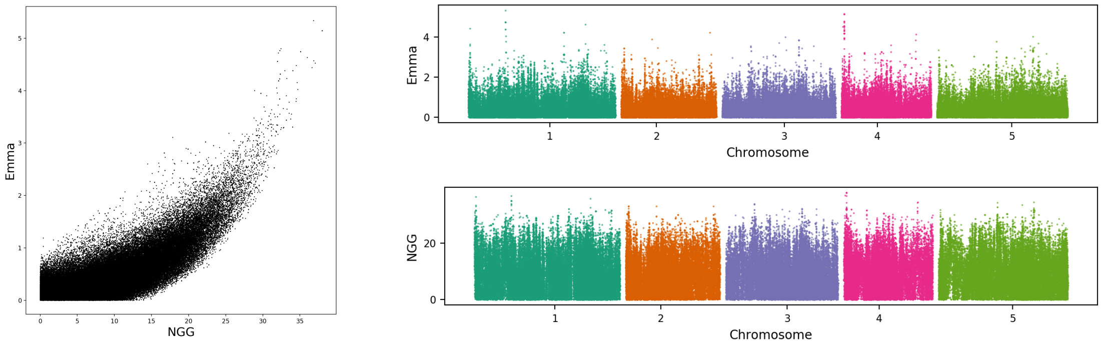


104
days to flowering trait
(SDV)

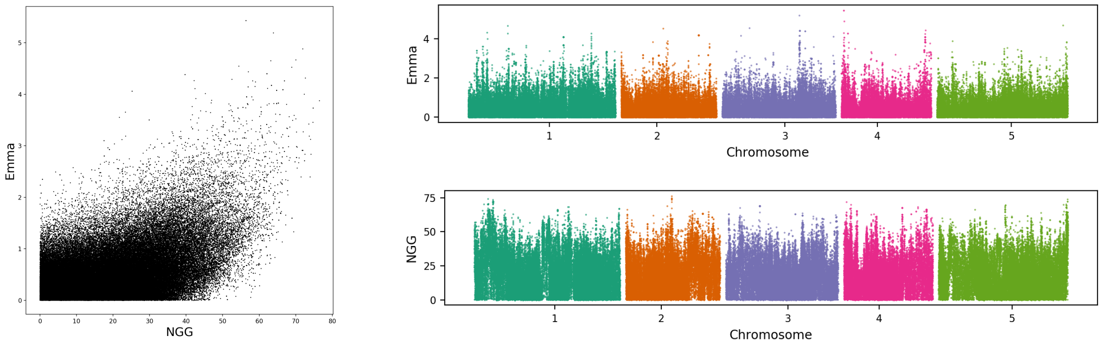


105
magnesium concentration
(Mg25)

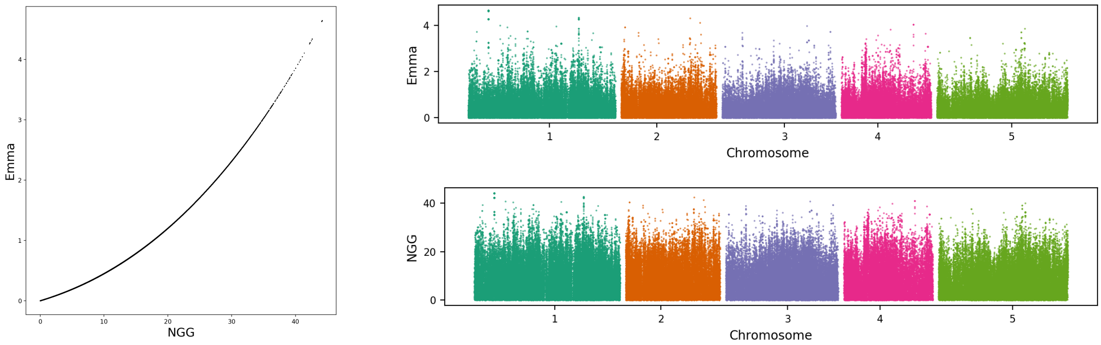


106
germination ratio
(Secondary Dormancy)

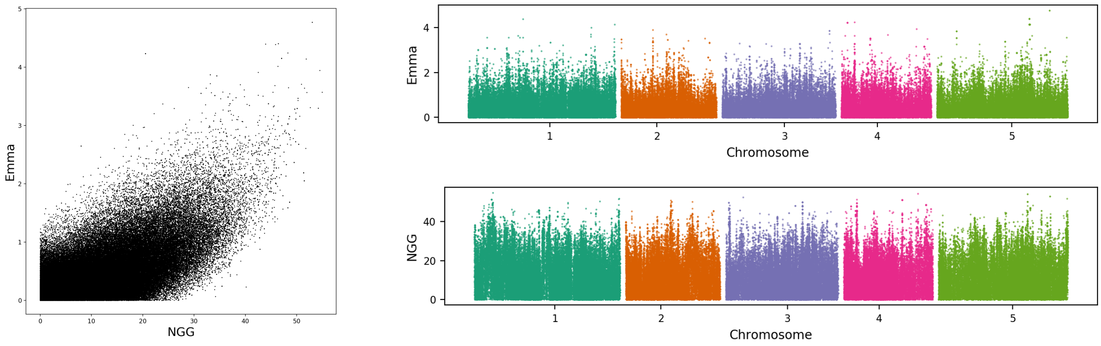

107
bacterial disease
resistance
(As)

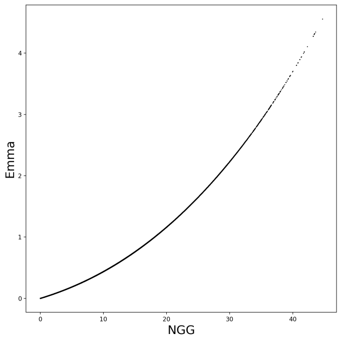


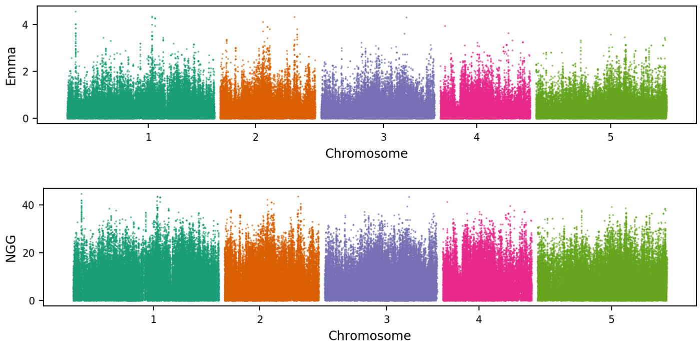
